# Interplay of Structural Heterogeneity and Active Remodeling Controls Chromatin Condensate Organization and Dynamics

**DOI:** 10.64898/2026.06.24.734404

**Authors:** Ritu Raj, Ranjith Padinhateeri, P. B. Sunil Kumar

## Abstract

Chromatin is an actively remodeled polymeric system whose organization emerges from the interplay of equilibrium interactions and ATP-dependent processes. Recent in vitro experiments show that nucleosome spacing and ATP-dependent remodeler activity significantly influence chromatin condensate properties. Here, guided by these observations, we develop a hierarchy of coarse-grained models that systematically dissect the roles of nucleosome spacing, remodeler-mediated binding–unbinding, and active force generation in governing condensate dynamics. We demonstrate that nucleosome spacing heterogeneity is a key determinant of condensate material properties. Condensates formed from regularly spaced fibers exhibit enhanced internal mixing, whereas those assembled from disordered spacing develop pronounced structural correlations, increased entanglement, and suppressed internal dynamics. Incorporating remodeler-like binding–unbinding non-equilibrium kinetics drives local structural reorganization, leading to condensate swelling and a substantial acceleration of internal relaxation. In a condensate of heterogeneous fibers, contrasts in spacing and activity robustly drive spatial segregation, giving rise to stable core–shell architectures. Strikingly, when dipolar forces are coupled to hydrodynamic interactions, serving as a minimal representation of active nucleosome translocation, condensates exhibit enhanced center-of-mass motion. Together, our results establish a predictive coarse-grained framework that quantitatively links structural heterogeneity and active processes to emergent chromatin-like condensate organization, mechanics, and transport.

**SIGNIFICANCE STATEMENT:** Chromatin forms dynamic condensates in living cells that play important roles in genome organization and regulation. Recent *in vitro* experiments have shown that condensate properties are strongly influenced by the arrangement of nucleosomes, the protein complexes around which DNA is wrapped, along chromatin fibers and by ATP-dependent remodeling activity. However, the mechanisms linking these molecular processes to condensate behavior remain poorly understood. Using a hierarchy of coarse-grained models, we identify the distinct roles of structural heterogeneity, remodeler-mediated kinetics, and active force generation in chromatin condensates. By providing a mechanistic explanation for experimental observations, our work establishes a physical framework connecting chromatin architecture and energy-consuming processes to condensate organization, dynamics, and function.

## I. INTRODUCTION

Genomic DNA in eukaryotic cells is packaged inside the nucleus in the form of chromatin, a hierarchical complex of DNA wrapped around histone proteins and associated regulatory factors [1]. The fundamental unit of chromatin is the nucleosome, in which ~147 base pairs (bp) of DNA are wrapped around a histone octamer [2, 3]. Nucleosomes slide and reposition along DNA, driving ATP-dependent chromatin remodeling [4, 5]. These regulated nucleosome movements organize DNA into arrays, controlling genome accessibility and influencing essential nuclear processes such as transcription, replication, and DNA repair [6, 7].

In *in vivo* systems, chromatin organization is inherently heterogeneous: its structure varies widely across the genome due to differences in linker DNA length, transcriptional activity, and local chromatin composition [8–11]. Additionally, ATP-dependent chromatin remodelers render this structural heterogeneity intrinsically time-dependent [4, 5]. These remodelers generate active forces on the chromatin fiber and surrounding medium, driving the system away from equilibrium and sustaining nonequilibrium steady states with distinct dynamical signatures [12–14].

Beyond local nucleosome repositioning, chromatin organization also exhibits strong spatial heterogeneity at the nuclear scale, with chromatin density varying across distinct genomic and nuclear regions [15]. Understanding how this density variability contributes to chromatin condensation has motivated extensive studies of chromatin-fiber organization [16, 17]. At a broad level, chromatin is commonly classified into highly compact heterochromatin and relatively open euchromatin regions. Recent studies have further shown that specific chromatin-associated regions can form condensates in vivo, including heterochromatin-rich domains and transcriptionally active condensates [18– 21].

To disentangle the contributions of different processes in such complex *in vivo* systems, several modeling and experimental approaches have investigated chromatin organization under simplified conditions [22–24]. In one class of models, chromatin is represented as a bead–spring polymer that undergoes condensation through effective intra-chromatin interactions, often mediated by chromatin-associated proteins [25]. In complementary regimes where stable condensate formation is not observed, other studies have focused on the three-dimensional organization of chromatin fibers, predicting both irregular fiber conformations and locally ordered short-range arrangements [26, 27].

To investigate the roles of nucleosome positioning along the fiber and nucleosome–nucleosome interactions, reconstituted chromatin has been used to generate condensates from short chromatin fibers [28–32]. Under physiological salt conditions, these fibers undergo liquid–liquid phase separation and form micron-sized condensates with nucleosome concentrations comparable to those observed *in vivo* [28, 32]. These experiments show that nucleosome spacing influences the material properties of chromatin condensates, with regularly spaced nucleosome arrays forming less viscous condensates than irregular arrays. In addition, remodeler activity drives condensate swelling and ATP-dependent motion. Together, these results provide a quantitative framework for probing how structural heterogeneity and active processes jointly regulate chromatin condensates.

Existing models of chromatin condensates primarily focus on equilibrium interactions, capturing phase separation and material properties as functions of inter-nucleosome interaction strength and nucleosome spacing along the fiber [33–36]. In contrast, most nonequilibrium chromatin-polymer models are highly coarse-grained, typically operating at mesoscopic to genomic length scales and describing dynamics over ~100 kb to a few Mb [37–40]. However, remodeler-driven activity occurs at the nucleosome scale, where enzymes reposition nucleosomes and modify nucleosome–nucleosome interactions. How such nucleosome-scale activity gives rise to larger-scale chromatin dynamics remains an open question. Another crucial aspect of active condensate dynamics is hydrodynamic interactions [37, 41], whereby the motion of one segment induces long-range fluid-mediated interactions that affect other parts of the fiber or condensate. How nucleosome-scale activity couples to hydrodynamic interactions, and how this coupling influences the collective dynamics of chromatin condensates, is still unclear. To address this gap, we develop coarse-grained models that explicitly account for nucleosome spacing and remodeler activity, and perform systematic simulations of nucleosome-scale chromatin fibers.

We present two coarse-grained models of chromatin condensates: a dry model evolved using Langevin dynamics and a hydrodynamic model incorporating explicit solvent via dissipative particle dynamics (DPD) [42, 43]. Although both operate at comparable coarse-grained resolution, they are constructed to isolate distinct physical mechanisms underlying condensate dynamics. The dry model incorporates a phenomenological switching mechanism that toggles nucleosomes between remodeler-bound and remodeler-unbound interaction states. In the hydrodynamic model, remodeler activity is additionally represented through the presence of stochastic active dipolar forces. A schematic overview of the two models is shown in Fig. 1.

**FIG. 1:**
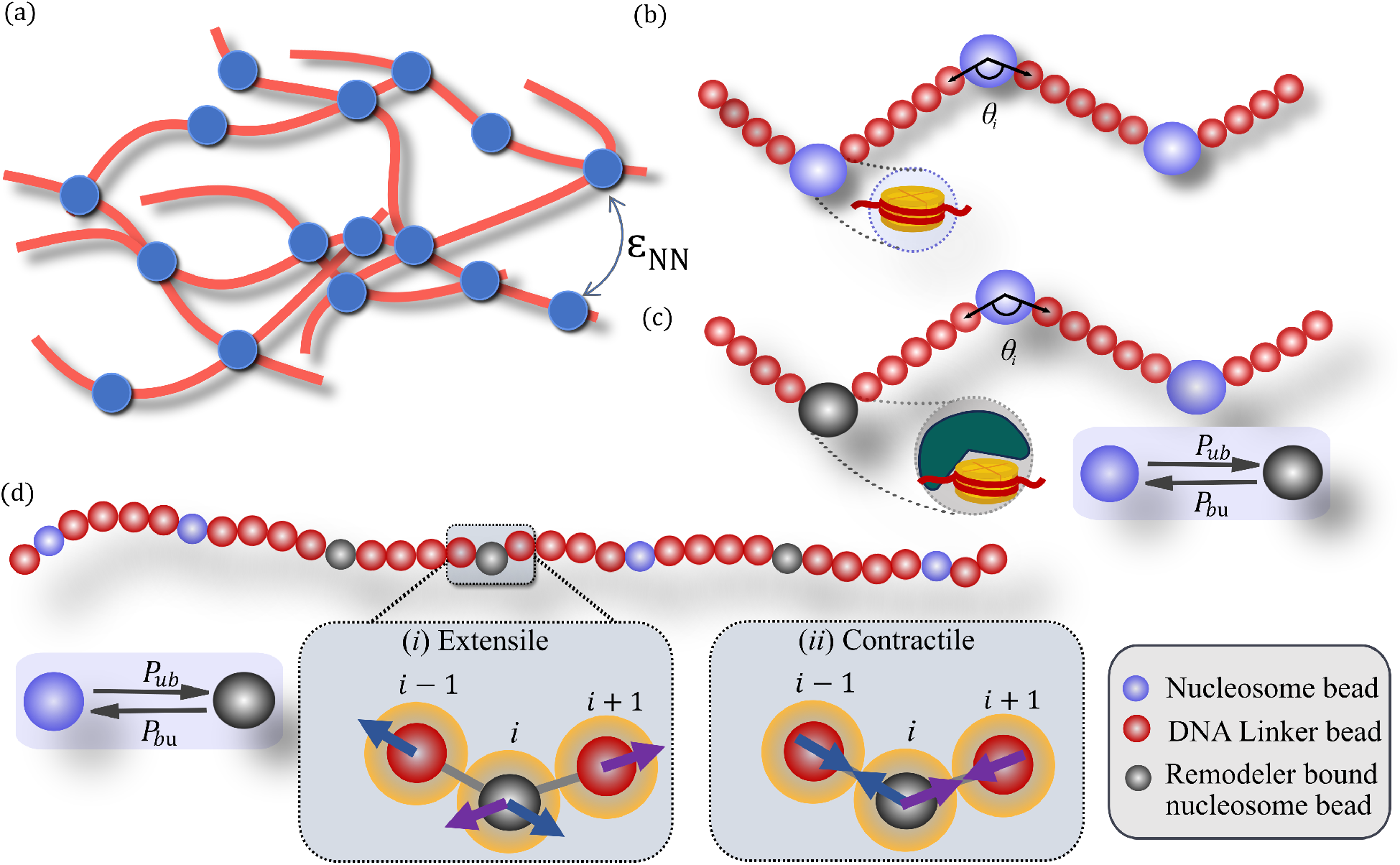
Coarse-grained chromatin fiber models incorporating ATP-dependent activity. **(a)** Schematic of multiple chromatin fibers interacting via inter-nucleosome attraction (*ϵ*_NN_), leading to condensate formation. **(b)** *Single-bead nucleosome model*. Nucleosomes are represented as beads of diameter 3*σ*connected by linker beads of diameter *σ*. The angle *θ*_*i*_ denotes the linker entry–exit angle at nucleosome *i*. **(c)** Nucleosomes stochastically switch between remodeler-unbound and remodeler-bound states with probabilities *P*_ub_ and *P*_bu_. **(d)** *Active chromatin model with hydrodynamics*. A nucleosome stochastically switches between remodeler-unbound and remodeler-bound states. In the bound state, dipolar forces are generated along the nucleosome–linker bonds corresponding to the dipole pairs (*i*− 1, *i*) and (*i, i*+1). These forces can be either *(i) extensile*, directed outward along the bond axis, or *(ii) contractile*, directed inward along the bond axis. The forces are distributed to solvent beads within spherical regions of radius *R*_*a*_ (shown in yellow), centered on the nucleosome bead *i* and its adjacent linker beads *j* (*j* = *i*± 1). Arrows indicate the resultant active force on the solvent within these spherical regions of radius *R*_*a*_. See Supplementary Methods (Sec. I) for details.

Our modeling framework reveals that condensates formed from uniformly spaced nucleosomes exhibit greater fluidity and hence faster internal mixing, whereas heterogeneous spacing leads to slower mixing and more constrained internal dynamics [28, 32]. Switching between remodeler-bound and unbound nucleosome states further enhances internal rearrangements and drives condensate swelling [32]. In mixed systems with fibers having differing nucleosome spacing or activity, we observe spontaneous segregation of fibers into a core–shell organization. Further, we find that incorporating stochastic active dipolar forces acting on nucleosome–linker segments enhances condensate motion [32].

Our modeling framework systematically disentangles the effects of structural remodeling and active force generation on chromatin condensates. The resulting picture distinguishes processes that fluidize and swell condensates from those that drive their motion, offering a physical framework for connecting remodeler activity to chromatin organization, accessibility, and regulation in both *in vitro* condensates and living cells.

## II. RESULTS

The self-organization and dynamics of multiple chromatin fibers are investigated in the presence and absence of nucleosome-remodeling activity. Simulations comprise *N*_*fiber*_ chromatin fibers resolved at the nucleosome level (Fig. 1a). Attractive inter-nucleosome interactions drive the spontaneous assembly of these fibers into a single condensate (Fig. S1b). The structural and dynamical properties of the resulting condensates are then characterized. Unless otherwise stated, all measurements are performed in the steady state, identified by the saturation of the condensate radius of gyration (see SI).

### 1. Inter-nucleosome Interaction Strength Controls Internal Dynamics in Chromatin Condensates

Passive condensate formation is first examined for chromatin fibers with a uniform linker length. Systematic variation of the inter-nucleosome interaction strength *ϵ*_NN_, reveals that condensates emerge for *ϵ*_NN_ ⪆ 3 *k*_B_*T* (Fig. S1). To assess condensate fluidity, the mean-squared displacement (MSD) of beads (representing nucleosome and linker), 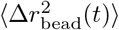, was calculated relative to the center of mass (CM) of the condensate. The MSD for different interaction strengths in the range 3 *k*_B_*T* ≤*ϵ*_NN_ ≤6 *k*_B_*T* is shown in Fig. 2a.

**FIG. 2:**
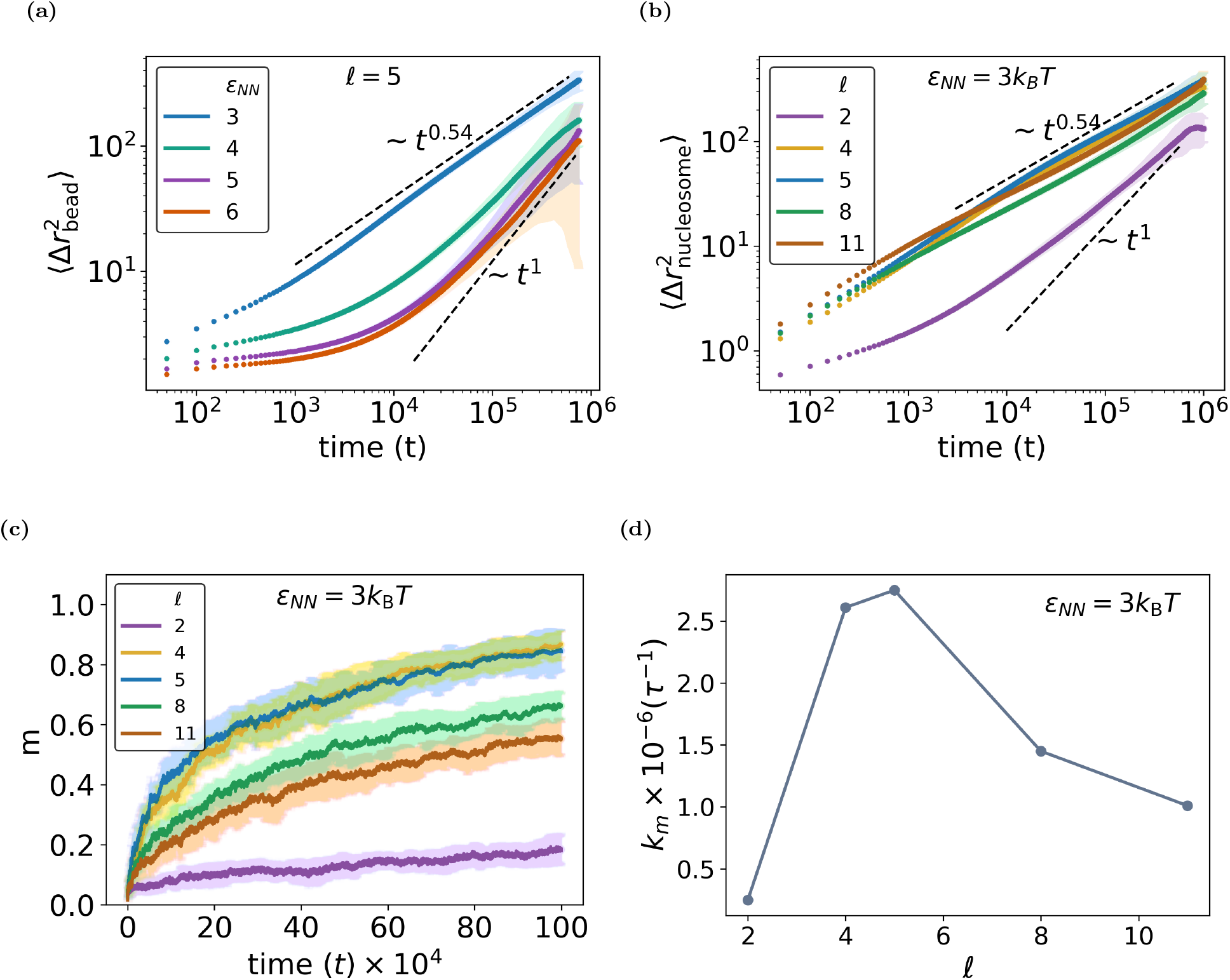
Effect of inter-nucleosome interaction strength *ϵ*_NN_ and linker length *ℓ* on chromatin condensate. **(a)** MSD of fiber beads measured in the center-of-mass frame of the condensate for different values of *ϵ*_NN_ at fixed linker length *ℓ* = 5. **(b)** MSD of nucleosome beads measured in the center-of-mass frame of the condensate for different linker lengths *ℓ* at fixed interaction strength *ϵ*_NN_ = 3*k*_B_*T*. **(c)** Mixing parameter *m*(*t*) as a function of time for different linker lengths *ℓ* at fixed *ϵ*_NN_ = 3*k*_B_*T*. **(d)** Mixing rate *k*_*m*_ corresponding to panel (c), obtained by fitting the mixing curves to 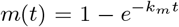, plotted as a function of linker length. The corresponding mixing timescale is defined as *t*_*m*_ = 1*/k*_*m*_. Shaded regions in **(a)–(c)** represent the standard deviation.

The generalized diffusion coefficient *D* and the dynamic exponent *α* were extracted by fitting the MSD, ⟨Δ*r*^2^(*t*)⟩ = 6*D t*^*α*^ (see Table S1). For all interaction strengths, at early times the MSD of the beads exhibits sub-diffusive behavior, with the extent of motion becoming increasingly restricted as *ϵ*_NN_ increases. This is shown in Fig. 2a for a linker length *ℓ* = 5. For *ϵ*_NN_ = 3 *k*_B_*T*, the dynamics exhibit a rapid crossover to a regime consistent with Rouse-like behavior, characterized by *α* ≈ 0.54 [40, 44]. As *ϵ*_NN_ is increased, a second crossover emerges at longer times, with the MSD approaching a diffusive scaling, *α* = 1 [45]. This behavior reflects the progressive collapse of individual fibers, after which the motion is dominated by the diffusion of their centers of mass. Consistent with this picture, the distribution of fiber *R*_*g*_ becomes narrower and shifts toward smaller values with increasing *ϵ*_NN_, indicative of enhanced compaction (Fig. S2a). The corresponding fiber center-of-mass MSD displays strongly constrained motion at short times, followed by a crossover to diffusion at longer times (Fig. S2b).

To relate the interaction strength used in the simulations to experimental chromatin-condensate dynamics, the characteristic bead displacement is compared with experimental measurements. FRAP experiments on reconstituted chromatin condensates report recovery over a timescale of ~5 min for condensates with radii *r* ≃4–5 *μ*m [28, 32]. Using the *D* and *α* obtained from the simulations, the characteristic displacement can be estimated as 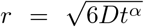, at *t* ≃ 5 min, for each *ϵ*_NN_ (Table S1). Among the interaction strengths considered, *ϵ*_NN_ = 3 *k*_B_*T* gives *r* ≃ 4.3 *μ*m, comparable to the experimental condensate radius. We therefore use *ϵ*_NN_ = 3 *k*_B_*T* as the inter-nucleosome interaction strength in the Langevin dynamics simulations, unless otherwise specified.

### 2. Linker Length Modulates Internal Dynamics and Mixing in Condensates

Given that the persistence length of DNA is approximately 150 bp, the spacing between neighboring nucleosomes is expected to play an important role in determining chromatin condensate properties. To investigate this effect, we consider fibers containing twelve nucleosomes separated by a uniform linker length *l*. The influence of *l* on both nucleosome-level and fiber-level dynamics within the condensate is then examined.

As shown in Fig. 2b, the MSD of nucleosomes exhibits Rouse-like behavior for linker lengths *ℓ >* 2, with the long-time scaling independent of linker length [40, 44]. However, for *ℓ* = 2, individual nucleosome motion is more constrained, reflecting tighter packing along the fiber. In this case, the MSD scales linearly with *t*, indicating that individual fibers collapse and move as single units [45]. This behavior is further confirmed by the corresponding MSD of the fiber CM, shown in Fig. S3.

The fluidity and internal mixing of beads within the condensate is further quantified using a mixing parameter *m*(*t*). To calculate *m*(*t*) the beads in one half of the condensate are tagged and the mixing of tagged and untagged beads are tracked over time (see Sec. I B 3 in the SI). Here, *m*(*t*) = 0 indicates complete separation between tagged and untagged beads, while *m*(*t*) = 1 indicates complete intermixing. The behavior of *m*(*t*) is shown in Fig. 2c for different linker lengths at a fixed *ϵ*_NN_. Fitting the data to 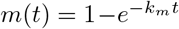, leads to a mixing timescale *t*_*m*_ = *k*^*−*1^. As shown in Fig. 2d, the mixing rate *k*_*m*_ varies non-monotonically with *ℓ*, increasing sharply from *ℓ* = 2 to *ℓ* = 4, peaking at *ℓ* ≈4–5, and then decreasing gradually, consistent with the corresponding trend in nucleosome MSD shown in Fig. 2b. The decrease in the mixing rate at larger linker lengths may arise from increased inter-fiber entanglements, which hinder bead rearrangements.

### 3. Irregular Nucleosome Spacing Decreases Fluidity of Chromatin Condensates

To examine the effect of nucleosome-spacing regularity on condensate properties, we compare condensates formed by regular- and irregular-fiber configurations. In the regular-fiber case, each fiber contains a fixed number of twelve nucleosomes with uniform spacing, *ℓ* = 5. In the irregular-fiber case, nucleosome spacing is chosen to follow experimentally observed distributions [32]. The experiments report variability in both nucleosome number and spacing, with each fiber displaying random spacing between nucleosomes [32]. To account for this variability in the simulations, the number of nucleosomes per fiber is drawn from a Gaussian distribution with mean 16 and standard deviation 2, truncated between 10 and 22 to exclude nonphysical extremes [32]. For each fiber in the simulation, the genomic length is fixed at 3.5 kbp [32], and the sampled nucleosomes are placed randomly along the fiber, resulting in an exponential linker-length distribution (Fig. S4).

To compare nucleosome mobility within the condensate, the MSD of nucleosomes relative to the condensate center of mass is analysed as a function of time (Fig. 3a). Both regular-fiber and irregular-fiber configurations exhibit 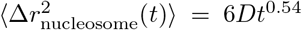.

**FIG. 3:**
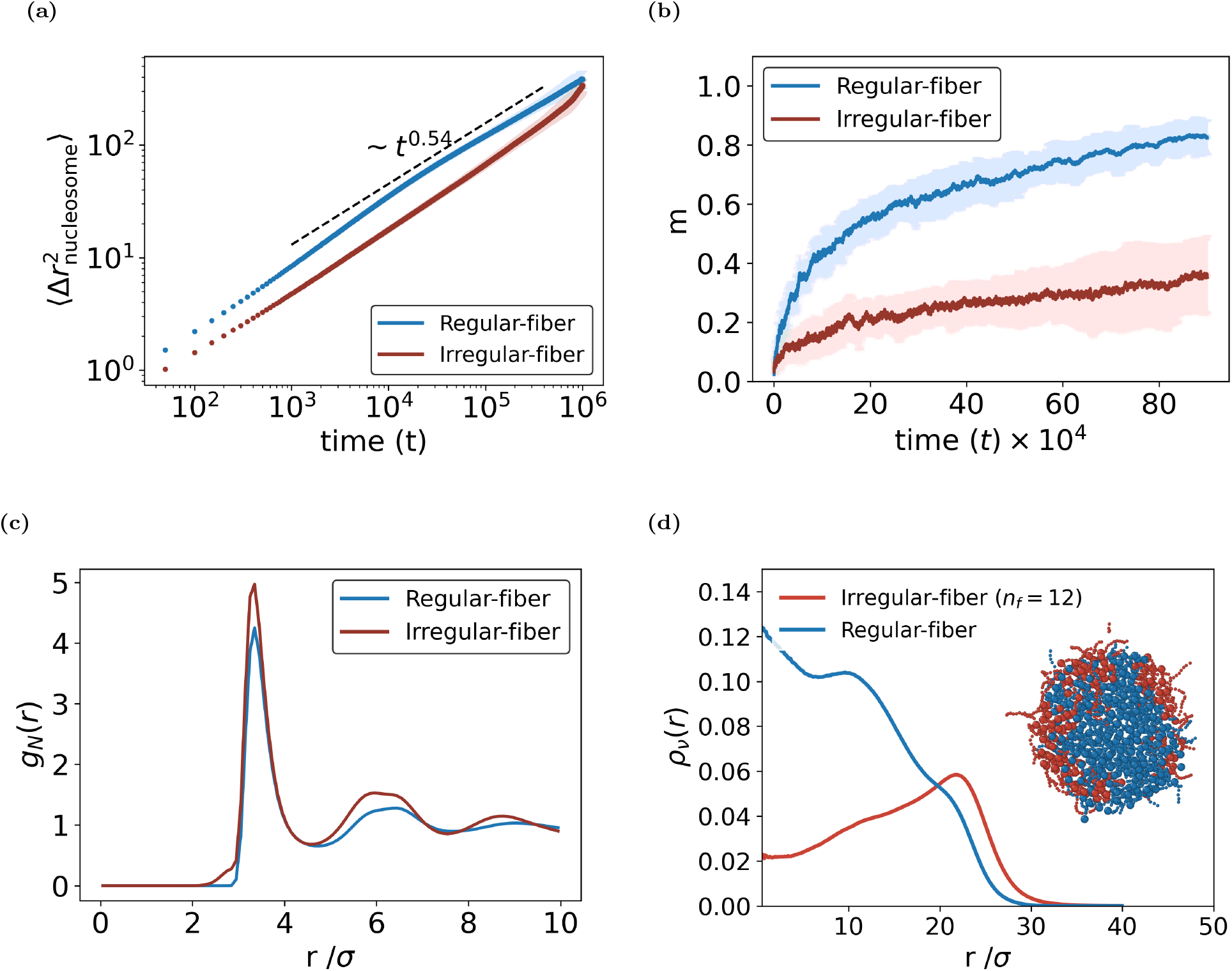
Comparison of condensate properties for regular- and irregular-fiber configurations. **(a)** MSD of nucleosomes measured in the condensate center-of-mass frame. **(b)** Mixing parameter *m*(*t*) as a function of time for the two configurations. **(c)** Nucleosome pair correlation function for the two configurations. **(d)** Radial bead density *ρ*_*ν*_ (*r*) of condensates formed from an equal mixture of regular (blue) and irregular (maroon) fibers, each containing *n*_*f*_ = 12 nucleosomes. *Inset:* Slab cut through the condensate corresponding to |*z* − *z*_CM_| ≤5. Shaded regions in (a) and (b) represent standard deviation.

However, the generalized diffusion coefficient *D* is significantly higher in the regular-fiber case (*D* = 0.04) compared to the irregular-fiber case (*D* = 0.01), indicating that nucleosomes in regular-fiber condensates exhibit higher mobility than those in irregular-fiber condensates.

FRAP experiments by Moore et al. [32] reported substantially slower fluorescence recovery in condensates composed of irregularly spaced nucleosomes compared to those with regular spacing. To quantify this behavior in simulations, we measure bead mixing using the parameter *m* (Eq. S21) for condensates formed from regular and irregular fiber configurations (see Sec. I B 3 in the SI). Beads in regular-fiber condensates exhibit efficient intermixing compared to irregular-fiber condensates, indicating reduced fluidity and a less dynamic internal structure in irregular fiber condensate (Fig. 3b).

The observed differences in nucleosome mobility and mixing may originate from differences in the underlying structural organization of nucleosomes within the condensate. To examine this possibility, the nucleosome pair correlation function (*g*_*N*_ (*r*)) was computed to characterize spatial correlations and local packing(Eq. S22) [46]. Condensates formed from irregular fibers exhibit substantially higher peaks in *g*_*N*_ (*r*) than those formed from regular fibers (Fig. 3c), indicating stronger local packing and more persistent spatial correlations. By contrast, regular-fiber condensates display weaker correlations and a more liquid-like organization.

Though the above comparison between regular and irregular fiber condensates was performed using irregular fibers in which the number of nucleosomes was sampled from a Gaussian distribution, analysis of irregular-fiber condensates with twelve randomly spaced nucleosomes per fiber showed only marginal changes in the results (Figs. S5a and S5b). In addition, the condensate center-of-mass MSD remains largely unaffected by nucleosome spacing, indicating that spacing heterogeneity mainly alters internal condensate dynamics rather than whole-condensate motion (Fig. S6).

To mimic ACF-remodeled chromatin fibers [32, 47], the number of nucleosomes per fiber is sampled from a Gaussian distribution, with nucleosomes equally spaced along each fiber. This produces fibers with regular intra-fiber organization but varying linker lengths across the population. The resulting condensate displays structural and dynamical characteristics that are much closer to the irregular-fiber case than to the regular-fiber case (Figs. S7a–S7c). The origin of this behavior lies in the linker-length distribution (Fig. S7d), which is enriched in longer linkers. These longer linkers promote entanglement and hinder nucleosome mixing, in agreement with the linker-length dependence shown in Fig. 2d.

### 4. Remodeler-driven Switching Induces Condensate Swelling and Enhances Internal Dynamics

We next investigate how remodeler-mediated binding–unbinding influences chromatin condensates. Nucleosomes stochastically transition between a remodeler-bound state, characterized by purely excluded-volume interactions, and a remodeler-unbound state, in which attractive inter-nucleosome interactions are present (see Model and Methods, Sec. I 1 in the SI and Fig. 1(c)). This switching serves as a minimal nonequilibrium model for the transient binding and dissociation of chromatin remodelers, dynamically regulating nucleosome contacts within the condensate. For simplicity, steric effects associated with the finite size of the remodelers are neglected.

To examine the effects of remodeler activity, we consider condensates formed from fibers with irregular nucleosome spacing and *n*_*f*_ = 12. The mean fraction of remodeler-bound nucleosomes, ⟨*n*_*b*_*/n*_*N*_⟩, is chosen to be 0.1 and 0.2, consistent with experimentally reported occupancies [32]. The switching dynamics are characterized by a switching time, *t*_a_, during which a fixed number, *n*_*b*_, of nucleosome state-switching attempts are performed (see Sec. I 1 in the SI). Importantly, switching is initiated only after the condensate has formed and the system has reached a steady state.

Introducing stochastic switching between remodeler-bound and remodeler-unbound nucleosome states leads to a pronounced swelling of the condensate. To quantify this effect, we calculate the condensate radius of gyration, *R*_*g*_ [48]. The variation of the time-averaged radius of gyration, ⟨ *R*_*g*_⟩, of the condensate with switching time, *t*_a_, is shown in Fig. 4a. A finite switching rate results in a systematic increase in *R*_*g*_. This behavior can be understood by comparing the switching timescale with the fiber relaxation time, 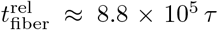, obtained from the autocorrelation of the unit end-to-end vector, 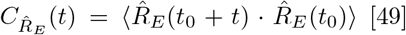. Since 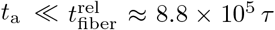, nucleosomes repeatedly change state before the surrounding chromatin can fully reorganize. In passive condensates, nucleosomes participating in attractive contacts (remodeler unbound nucleosomes) are preferentially located in the condensate interior, whereas linker segments and weakly interacting regions (remodeler bound nucleosomes) are enriched near the interface. State switching continuously disrupts and reforms these contacts, driving nucleosomes between interior and interfacial environments. Because the chromatin fibers cannot fully relax between successive switching events, the system samples increasingly expanded conformations. The distributions of remodeler bound and unbound nucleosomes in passive and active condensates, shown in Figs. S8 and S9, are consistent with this scenario. The resulting increase in configurational entropy competes with the attractive nucleosome–nucleosome interactions that stabilize the condensate, leading to a larger steady-state condensate size. Consistent with this mechanism, the effect is stronger for ⟨*n*_*b*_*/n*_*N*_⟩ = 0.2, where a larger fraction of remodeler-bound nucleosomes generates more frequent contact rearrangements.

**FIG. 4:**
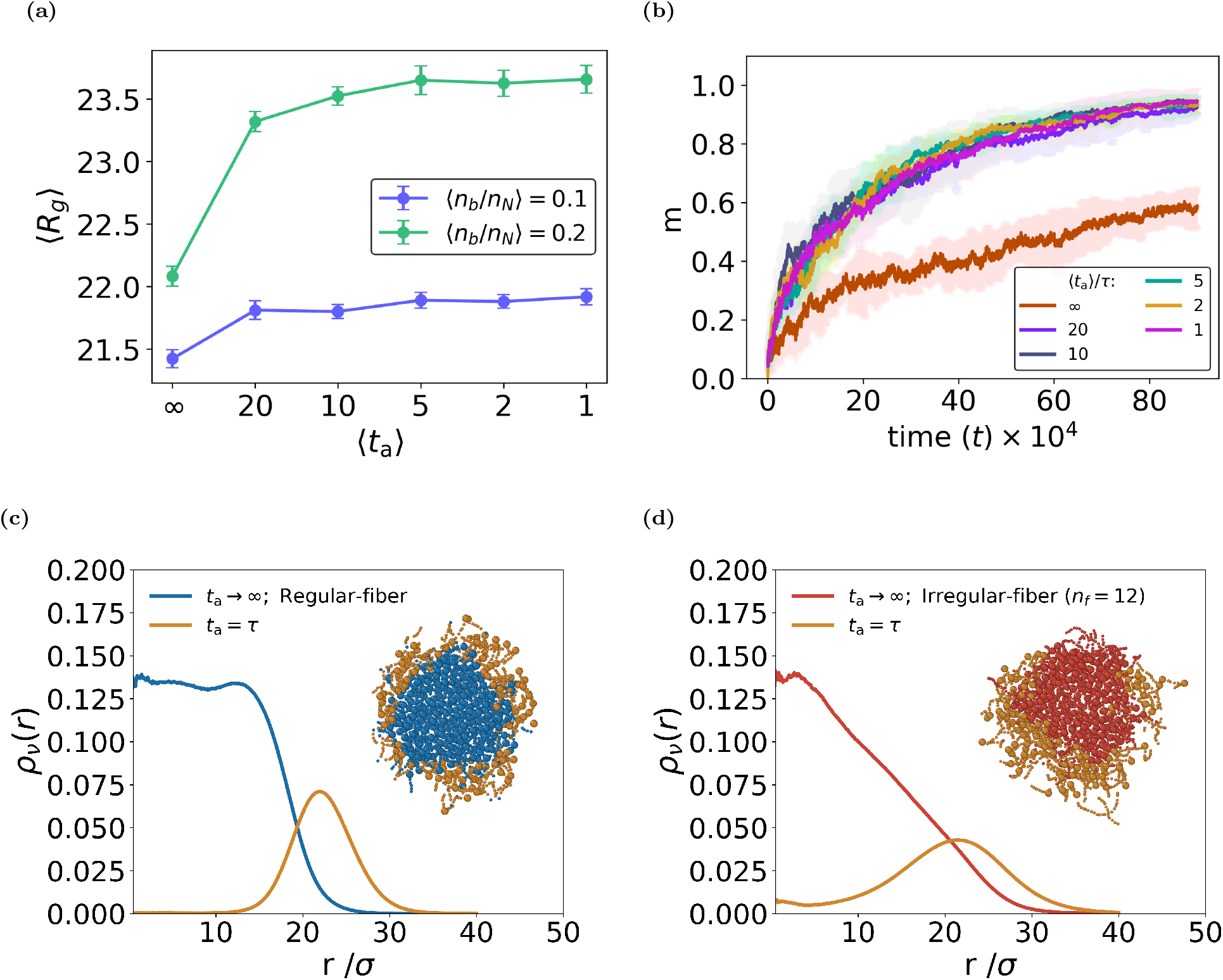
Effect of remodeler binding–unbinding activity on a chromatin condensate. **(a)** Radius of gyration (*R*_*g*_) of the condensate as a function of the switching time *t*_a_, for different average fractions of remodeler-bound nucleosomes ⟨*n*_*b*_*/n*_*N*_⟩. **(b)** Mixing parameter as a function of time for the same systems as in (a) at fixed ⟨*n*_*b*_*/n*_*N*_⟩ = 0.1. Shaded regions represent the standard deviation. **(c**,**d)** Radial bead density profiles of a condensate composed of an equal mixture switching-active fibers and either regular-fiber (c) or irregular-fiber with a fixed number of twelve nucleosomes per fiber (d), shown for *t*_a_ = *τ* and ⟨*n*_*b*_*/n*_*N*_⟩ = 0.1. *Inset*: Slab cut through the condensate corresponding to | *z* − *z*_CM_|≤ 5; regular fibers: blue, irregular fibers: maroon, and switching-active fibers: golden yellow.

The continual rearrangement of nucleosome contacts induced by remodeler binding–unbinding dynamics also enhances transport within the condensate. To quantify this effect, we compute the bead mixing parameter (Eq. S21) for different switching times, *t*_a_. As shown in Fig. 4b and Fig. S10a, the introduction of switching leads to a substantial increase in mixing, indicating enhanced internal mobility. The extent of mixing further increases with the average fraction of remodeler-bound nucleosomes, ⟨*n*_*b*_*/n*_*N*_⟩, consistent with greater condensate fluidisation at higher remodeler occupancies. This enhanced mixing is accompanied by increased fiber motion within the condensate, as evidenced by the mean-squared displacement of fiber centers of mass relative to the condensate center of mass (Fig. S11a, Fig. S10b). Interestingly, once switching is activated, both the mixing and the fiber MSD exhibit only a weak dependence on the switching time, *t*_a_. In contrast, the center-of-mass motion of the condensate itself remains essentially unchanged, as demonstrated by the condensate CM MSD (Fig. S11b). These results indicate that remodeler-mediated switching primarily fluidises the condensate and accelerates internal reorganization, while leaving the overall translational motion of the condensate unaffected.

Taken together, these findings demonstrate that remodeler-driven switching leads to condensate swelling and substantially enhances internal dynamics, increasing fiber mobility and mixing, whereas the translational motion of the condensate center of mass remains essentially unchanged.

### 5. Spatial Segregation in Condensates Driven by Differences in Nucleosome Spacing and Remodeler

*Activity*

Having characterized condensates formed from homogeneous fiber populations, we next investigate condensates assembled from mixtures of chromatin fibers under three distinct conditions: (i) mixtures of passive regular and irregular fibers (*t*_a_→∞), (ii) switching-active fibers (*t*_a_ = *τ*) mixed with passive regular fibers, and (iii) switching-active fibers (*t*_a_ = *τ*) mixed with passive irregular fibers. In each case, the condensate comprises a total of 108 fibers with equal numbers of each fiber type, and each fiber contains twelve nucleosomes. The average fraction of remodeler-bound nucleosomes was fixed at ⟨*n*_*b*_*/n*_*N*_ ⟩ = 0.1, with switching time *t*_a_ = *τ*.

In all three cases, the mixed condensates undergo spatial segregation, giving rise to a stable core–shell structure in which one fiber population is concentrated in the condensate interior, while the other is enriched near the surface (see slab cross-sections in the insets of Figs. 3d, 4c, and 4d).

To quantify the spatial seggregation, we compute the radial bead density *ρ*_*ν*_ (*r*) for each fiber population *ν* (Eqn. S24). In all three systems, the radial density profiles reveal a clear core–shell organization. For mixtures of regular and irregular fibers [case (i)], regular fibers preferentially occupy the condensate interior, while irregular fibers are enriched near the surface (Fig. 3d). For mixtures of passive and switching-active fibers [cases (ii) and (iii)], passive fibers form the core and switching-active fibers are preferentially localized at the periphery (Figs. 4c and 4d).

The segregation observed in all three systems appears to be governed by differences in the equilibrium packing density of the constituent fibers. Fiber populations that favor denser packing preferentially occupy the condensate interior, while populations that form more expanded, lower-density structures are enriched near the interface. Since irregular linker spacing and remodeler-mediated switching both promote swollen, less densely packed conformations, irregular and switching-active fibers are consistently expelled from the condensate core, resulting in the observed core–shell organization.

### 6. Active Forces and Hydrodynamic Interactions Enhances Chromatin Condensate Motion

Experiments indicate that ATP-dependent chromatin remodelers enhance the translational motion of chromatin condensates, with the magnitude of motion increasing with ATP concentration [32]. Such ATP-dependent effects can arise from two distinct contributions: (a) remodeler binding–unbinding, which alters nucleosome interactions, and (b) active forces generated by remodelers on the chromatin and the surrounding medium during nucleosome translocation. As shown earlier, when remodeler-induced interaction switching is incorporated within a Langevin framework without hydrodynamic interactions, no appreciable change in the overall condensate motion is observed, and the center-of-mass dynamics remain indistinguishable from those of the passive system (Fig. S11b).

ATP-dependent condensate motility is hypothesized to arise from active forces generated during nucleosome translocation and their hydrodynamic coupling to the solvent. To capture this mechanism, dissipative particle dynamics (DPD) simulations are performed with active force dipoles of strength *F*_0_. These force dipoles represent the leading-order active forcing consistent with local momentum conservation. Details of this implementation are provided in the *Model and Methods* section of the SI, Sec. I 2.

With the DPD simulations, we first examine the monomer dynamics within the condensate in the absence of active force (*F*_0_ = 0). The MSD of beads in the condensate center-of-mass frame exhibits the expected Zimm scaling, ~*t*^2*/*3^, for both with and without nucleosome state switching (Fig. 6a) [44], confirming the presence of hydrodynamic coupling. Additionally, we observe that hydrodynamic interactions do not alter the enhanced dynamics resulting from nucleosome state switching.

A clear enhancement in condensate motion is observed on introduction of active forcing, *F*_0_ ≠ 0 (Fig. 5), as can be seen in the condensate center-of-mass MSD for extensile dipolar forces *F*_0_ *>* 0 (Fig. 6b; see also Movie S1). At long times, the MSD exhibits diffusive behavior for all values of *F*_0_, with a linear scaling 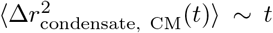, indicating that dipolar active forcing primarily renormalizes the effective diffusion coefficient without altering the transport exponent.

**FIG. 5:**
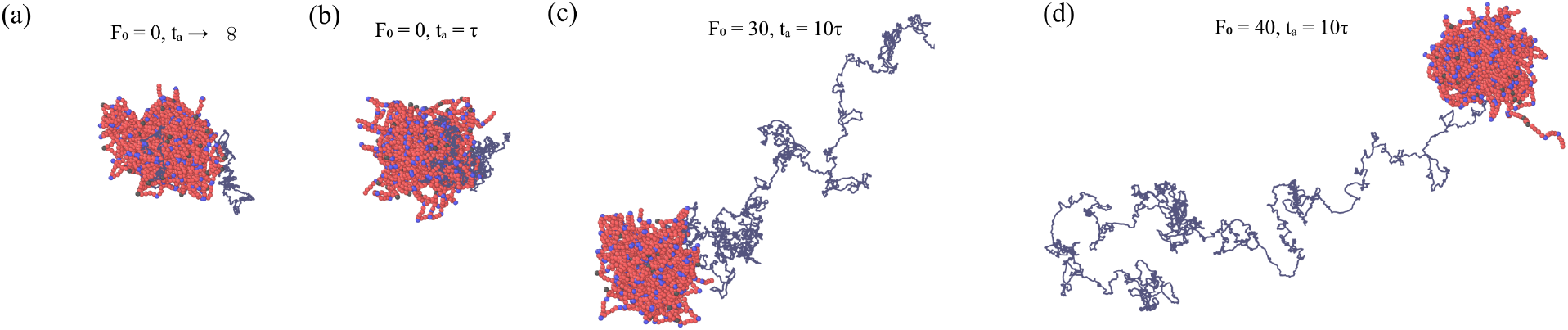
Snapshots of condensate center-of-mass trajectories under different activity conditions. The grey lines show the paths traced by the condensate center of mass over a fixed duration of 10^5^*τ*. **(a)** Passive case (*F*_0_ = 0, *t*_a_→ ∞). (b)Nucleosome state switching without dipolar activity (*F*_0_ = 0, *t*_a_ = *τ*). **(c**,**d)** Dipolar activity with *F*_0_ = 30 and 40, respectively. See Supplementary Movie S1.

**FIG. 6:**
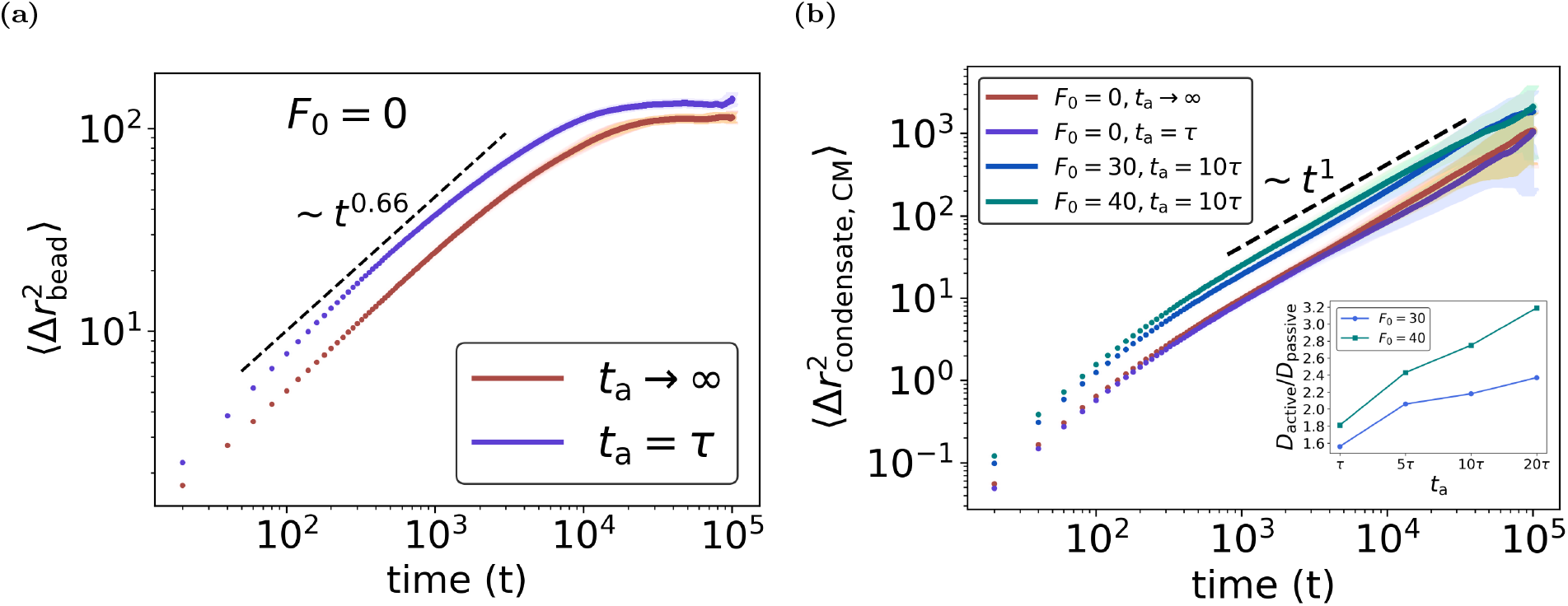
Active Chromatin Model with Hydrodynamics. **(a)** MSD of beads relative to the condensate CM for *F*_0_ = 0 and *t*_a_ = *τ* and ∞. **(b)** Condensate CM MSD for different values of *F*_0_ and *t*_a_. The passive case corresponds to *F*_0_ = 0, *t*_a_ → ∞. Inset: Diffusion-coefficient ratio, *D*_active_*/D*_passive_, as a function of *t*_a_ for *F*_0_ = 30 and 40. Shaded regions represent the standard deviation.

Additionally, for a fixed activity strength *F*_0_, the enhancement in the condensate center-of-mass MSD increases with increasing switching time *t*_a_, because a larger *t*_a_ increases the persistence time of the dipolar force acting on remodeler-bound nucleosomes and their bonded linkers (Figs. S12a, S12b).

This enhancement is characterized by the ratio *D*_active_*/D*_passive_, plotted as a function of *t*_a_ (inset of Fig. 6b). The passive condensate corresponds to *F*_0_ = 0 and *t*_a_→ ∞. For persistent activity, the diffusion coefficient increases by factors of approximately 2.4 and 3.2 relative to the passive case for *F*_0_ = 30 and 40, respectively. This enhancement is consistent with experimentally observed increases in condensate diffusivity, which are of comparable order of magnitude [32] (Fig. S13).

Further, the above-mentioned features remain largely unchanged for contractile dipolar forces (*F*_0_ *<* 0), closely matching those obtained for extensile dipolar forces. This indicates a weak dependence of the condensate dynamics on the sign of the dipolar activity (Fig. S14; see also Fig. S15 and Movie S2).

These results show that remodeler generated active forces, mediated through hydrodynamic interactions, drive the observed increase in condensate transport.

## III. DISCUSSION

Chromatin in the nucleus exhibits substantial heterogeneity in nucleosome spacing and remodeling activity [4, 8, 9, 50]. Recent *in vitro* studies have sought to understand the consequences of this heterogeneity by reconstituting chromatin condensates with controlled nucleosome spacing and remodeler activity[28–32]. These experiments have revealed that both factors strongly influence condensate structure and dynamics. However, because their effects are often intertwined, isolating their individual physical contributions remains challenging.

In this work, we employed a hierarchy of coarse-grained chromatin models to systematically isolate the effects of (i) nucleosome spacing, (ii) transient remodeler binding–unbinding that modulates internucleosome interactions, and (iii) active force generation by remodelers and its coupling to the surrounding solvent. By comparing these models, we disentangle the distinct physical roles of each mechanism and establish a unified framework that provides a consistent physical explanation for several experimental observations on *in vitro* chromatin condensates.

Our results demonstrate that, in the absence of active processes, nucleosome spacing is a key determinant of chromatin condensate properties. In particular, spacing heterogeneity tunes the rheological behavior of the condensate: regular-fiber arrays form more fluid assemblies, whereas irregular-fiber arrays produce condensates with higher viscosity and slower internal mixing, consistent with experimental observations [32].

The slower mixing dynamics in irregular-fiber condensates point to the existence of additional constraints on fiber rearrangements. A natural candidate is topological entanglement between chromatin fibers. To test this possibility, we quantify inter-fiber entanglement using the distribution of the absolute Gaussian linking number, | *Lk*_*G*_ |, computed for all fiber pairs (Fig. S16) [51–53] (see SI, Sec. I B 7).

Condensates formed from irregular fibers exhibit a broader distribution of |*Lk*_*G*_ | with significant weight at large linking numbers, reflecting higher topological entanglement relative to regular-fiber systems. The broad distribution of linker lengths, particularly the presence of long linker segments, in irregular fibers, increases the likelihood of inter-fiber interpenetration and topological trapping. Consequently, fiber motion becomes increasingly constrained, hindering structural rearrangements within the condensate and resulting in reduced nucleosome mobility and slower internal mixing. Notably, condensates with intra-fiber regular spacing, mimicking ACF-remodeled fibers, exhibit a similar distribution of |*Lk*_*G*_ | as the irregular-fiber case, indicating comparable levels of inter-fiber entanglement (Fig. S16). Consistent with this, these condensates display dynamics similar to those of irregular-fiber condensates [32].

The influence of entanglements is also reflected in nucleosome dynamics within the condensate. Although nucleosome motion remains subdiffusive, with ⟨Δ*r*^2^(*t*) ⟩ ~*t*^0.54^, consistent with Rouse-like dynamics, the effective diffusion coefficient (*D*_*N*_) is significantly lower for fibers with heterogeneous linker spacing than for uniformly spaced fibers, indicating slower nucleosome mobility. These results suggest that nucleosome spacing heterogeneity influences chromatin dynamics not only through its effect on local packing but also by modulating the degree of topological entanglement within chromatin assemblies. Consequently, regions with different linker-length distributions are expected to exhibit distinct dynamical behaviors [54–56]. Given the substantial variability in nucleosome spacing across the genome, entanglements associated with longer linker segments may therefore contribute to the regulation of chromatin accessibility and higher-order chromatin organization [33, 57].

Transient remodeler binding and unbinding to nucleosomes provide an additional source of internal dynamics by intermittently weakening inter-nucleosome attractions [4, 5]. In the simulations, switching was introduced after condensate formation to probe its effect on already formed structures. Introducing remodeler binding–unbinding dynamics prior to condensate formation suppresses phase separation when the switching timescale is shorter than the fiber relaxation time, highlighting the competition between nucleosome interaction lifetime and nucleosome mobility in stabilizing the condensed phase. This is consistent with experiments showing that remodeler activity can suppress or delay condensate formation [32].

Remodeler binding–unbinding dynamics enhance condensate swelling and accelerate internal mixing, in qualitative agreement with experimental observations [32]. These results demonstrate that modulation of inter-nucleosome interactions alone is sufficient to drive condensate expansion in *in vitro* systems. However, despite significantly altering condensate structure and internal dynamics, binding–unbinding activity does not enhance condensate center-of-mass motion. This indicates that remodeler-mediated contact remodeling primarily regulates condensate cohesion and fluidity, but does not inject momentum into the system or account for the experimentally observed ATP-dependent condensate motility.

A qualitatively different behavior emerges when active dipolar forces are incorporated, representing remodeler-activity and its hydrodynamic coupling. This activity, while preserving the long time diffusive motion of the condensate center of mass, enhances the effective diffusion coefficient. The enhancement depends on the dipole strength and activity duration, but only weakly on its sign, indicating that both extensile and contractile forces promote motion of the condensate.

Our results provides plausible explanation for experiments showing remodeler-specific effects on condensate motility; for example, RSC enhances motion whereas ACF does not [32]. Such differences may reflect variations in the magnitude or persistence of the active forces.

For condensates formed from mixed fiber populations, the observed core–shell organization arises from a competition between conformational entropy and effective inter-nucleosome cohesion. Such physical segregation of fibers with distinct structural properties suggests a possible mechanism that could contribute to the emergence of chromatin microdomains with distinct material characteristics within the nucleus [58].

We emphasize that this minimal coarse-grained framework is designed to isolate the roles of nucleosome spacing, remodeler-mediated binding– unbinding, and active force generation. Salt-dependent effects such as electrostatic screening, ion-mediated bridging, and multivalent-ion-specific interactions are not explicitly resolved in the present model; instead, chromatin compaction is controlled phenomenologically through an effective nucleosome–nucleosome interaction strength. Because the model resolves chromatin at the nucleosome scale, the accessible simulation times remain shorter than the much longer experimental timescales. Accordingly, the switching time and active-force duration should be interpreted as phenomenological parameters rather than quantities directly mapped to ATP concentration or ATP hydrolysis rates.

Within this framework, our results reveal that different chromatin remodeling related processes influence condensate behavior in distinct ways: spacing heterogeneity primarily controls material properties, remodeler binding–unbinding reorganizes internal structure, and active forces generated by remodeler drive condensate motion. By isolating these contributions within a unified framework, we provide a coherent physical picture of how structural, interaction, and active processes combine to govern chromatin condensate dynamics at the mesoscale.

## ACKNOWLEDGMENTS

We thank Geeta J. Narlikar, Vijay Ramani, and Camille Moore for valuable discussions regarding the experimental findings. Computational support from the PARAM Shakti supercomputing facility at IIT Madras is gratefully acknowledged. PBSK acknowledges the financial support provided by Anusandhan National Research Foundation (ANRF) under J. C. Bose grant (ANRF/JBG/2025/000187/PS).

