## Supplementary Information for "Interplay of Structural Heterogeneity and Active Remodeling Controls Chromatin Condensate Organization and Dynamics"

### I. MODELS AND METHODS

We investigate the effects of nucleosome arrangement and remodeler-induced activity on chromatin condensates using two coarse-grained models. The first is a Single-bead nucleosome model that captures the role of nucleosome arrangement and stochastic remodeler binding–unbinding. The second is an active chromatin model with hydrodynamic interactions, which incorporates remodeler-generated forces and their coupling to the surrounding solvent. A detailed description of the models is provided below.

Chromatin filaments are modeled as block cofibers composed of nucleosomes and semi-flexible linker DNA segments. The system contains $N_{fiber}$ fibers, with the $j^{th}$ fiber consisting of $N_{f,j}$ beads. The position of the $i^{th}$ bead is denoted by $\vec{r}_i(t)$, where $i = 1, \ldots, n$ and $n = \sum_{j=1}^{N_{fiber}} N_{f,j}$. We first describe the single-bead nucleosome model.

#### A. Single-bead nucleosome model

In the single-bead nucleosome model, a nucleosome (N) is represented by a bead of diameter $3\sigma$, while linker DNA is represented by beads of diameter $\sigma$ (Fig. 1b). The position $\vec{r}_i(t)$ of bead $i$ evolves according to the Langevin equation

$$m_i \frac{d^2\vec{r}_i(t)}{dt^2} = -\gamma \frac{d\vec{r}_i(t)}{dt} - \nabla_i U + \sqrt{2\gamma k_B T}\vec{\xi}_i(t). \quad \text{(S1)}$$

Here, $m_i$ is the bead mass and $\gamma$ is the friction coefficient. The term $-\nabla_i U$ denotes the conservative force acting on bead $i$, arising from bonded, non-bonded, and bending interactions defined below. The stochastic term $\vec{\xi}_i(t)$ is a Gaussian white noise satisfying $\langle \xi_{i\alpha}(t) \rangle = 0$ and $\langle \xi_{i\alpha}(t)\xi_{j\beta}(t') \rangle = \delta_{ij}\delta_{\alpha\beta}\delta(t - t')$, where $\alpha, \beta \in \{x, y, z\}$ denote Cartesian components.

Lengths, energies, and times are expressed in units of bead diameter $\sigma$, $k_B T$, and $\tau$, respectively, where $\tau = \sigma\sqrt{m/k_B T}$. In this coarse-grained mapping, $\sigma$ corresponds to 10 base pairs (bp) ($\approx$ 3.4 nm), ensuring that the contour length of the fiber and the number of nucleosomes match experimental systems [1, 2]. The bead mass is estimated by assuming a solvent density equal to that of water, yielding a physical time scale $\tau \approx 2 \times 10^{-10}$s. The friction coefficient is set to $\gamma = 1$, ensuring that inertial effects are negligible and the dynamics are effectively overdamped.

The potential energy $U$ is given by

$$U = \sum_{\langle i,j \rangle} U^{b}(r_{ij}) + \sum_{i<j} U^{nb}(r_{ij}) + \sum_{\langle i,j,k \rangle} U^{bend}(\theta_{ijk}). \quad \text{(S2)}$$

The term $U^{b}(r_{ij})$ represents the bonded interaction between neighboring beads $i$ and $j$, modeled as a harmonic spring:

$$U^{b}(r_{ij}) = \frac{1}{2} k_b (r_{ij} - r_0)^2, \quad \text{(S3)}$$

with spring constant $k_b = 20$. Here, $r_{ij}$ is the distance between beads $i$ and $j$, and $r_0$ is the equilibrium bond length, set to 1 for linker–linker bonds, 2 for linker–nucleosome bonds, and 3 for nucleosome–nucleosome bonds.

The term $U^{nb}(r_{ij})$ denotes the non-bonded interaction between beads $i$ and $j$, given by

$$U^{nb}(r_{ij}) = \begin{cases} u(r_{ij}) - u(r_c), & r_{ij} \le r_c, \\ 0, & \text{otherwise.} \end{cases} \quad \text{(S4)}$$

The functional form of $u(r_{ij})$ depends on the bead types involved. For nucleosome–nucleosome pairs, $u(r_{ij})$ is given by the Lennard–Jones (LJ) potential

$$u_{LJ}(r_{ij}) = 4\epsilon_{NN} \left[ \left( \frac{3\sigma}{r_{ij}} \right)^{12} - \left( \frac{3\sigma}{r_{ij}} \right)^{6} \right], \quad \text{(S5)}$$

*
†

with cutoff $r_c = 7.5$. For all other non-bonded pairs (i.e., linker–linker and linker–nucleosome interactions), $u(r_{ij})$ is given by the Weeks–Chandler–Andersen (WCA) potential,

$$u_{\rm WCA}(r_{ij}) = 4\epsilon_{\rm WCA}\left[\left(\frac{\sigma'}{r_{ij}}\right)^{12} - \left(\frac{\sigma'}{r_{ij}}\right)^{6} + \frac{1}{4}\right], \tag{S6}$$

where $\sigma' = 1$ for linker-linker and $\sigma' = 2$ for linker-nucleosome interactions. We set $\epsilon_{\rm WCA} = k_{\rm B}T$ with $r_c = 2^{1/6}\sigma'$.

The term $U^{\rm bend}(\theta_{ijk})$ represents the three-body bending interaction between consecutive beads $(i, j, k)$, which penalizes deviations from a preferred bond angle. It is given by

$$U^{\rm bend}(\theta_{ijk}) = \frac{1}{2}\kappa\left[1 - \cos\left(\theta_{ijk} - \theta_0\right)\right], \tag{S7}$$

where $\theta_{ijk}$ is the angle formed by the bonds $(i, j)$ and $(j, k)$ (see Fig. 1a), $\theta_0$ is the preferred bond angle, and $\kappa$ is the bending stiffness.

Unless otherwise specified, we set $\kappa = 28$, $\theta_0 = 180°$ for linker–linker–linker triplets, and $\kappa = 3.8$, $\theta_0 = 60°$ for linker–nucleosome–linker triplets [3, 4]. These constraints ensure a persistence length of approximately 140 bp for linker DNA [5, 6] and a preferred entry–exit angle for the linker–nucleosome–linker configuration [4].

*Binding–Unbinding Remodeler Activity*

To capture the effect of chromatin remodelers that transiently bind to nucleosomes and modulate inter-nucleosomal interactions, we incorporate stochastic switching between two states: a *remodeler-unbound* state, where nucleosomes interact via a LJ potential (Eq. S5), and a *remodeler-bound* state, representing the screened interaction state induced by remodeler binding and resulting in purely excluded-volume interactions. This is implemented by replacing the LJ interaction between a remodeler-bound nucleosome and all other nucleosomes with a WCA potential (Eq. S6, with $\sigma' = 3$ and $r_c = 2^{1/6}(\sigma')$). Remodelers are modeled implicitly via stochastic switching between nucleosome states, modulating effective nucleosome–nucleosome interactions (see Fig. 1c).

Nucleosome state transitions are executed stochastically. At each time interval $\tau$, $n_{\rm sw}$ nucleosomes are selected uniformly at random from the population of $n_N$ nucleosomes, and a state-switching attempt is performed for each selected nucleosome. If the selected nucleosome is in the remodeler-bound state, it switches to the unbound state with probability $\min(1, P_{\rm bu})$, where

$$P_{\rm bu} = \frac{1}{1 + \exp\left[\mu(n_{ub} - n_b - A_0)\right]}, \tag{S8}$$

and if the selected nucleosome is in the remodeler-unbound state, it switches to the bound state with probability $\min(1, P_{\rm ub})$, where

$$P_{\rm ub} = \frac{1}{\eta + \exp\left[-\mu(n_{ub} - n_b - A_0)\right]}. \tag{S9}$$

Here, $n_{ub}$ and $n_b$ denote the instantaneous numbers of remodeler-unbound and remodeler-bound nucleosomes, respectively, while $n^0_{ub}$ and $n^0_b$ represent their steady-state values. The parameter $\mu$ controls the magnitude of fluctuations in $n_{ub}$ and $n_b$. The term $A_0 = n^0_{ub} - n^0_b$ defines the asymmetry parameter, and $\eta = 2\left(\frac{n^0_{ub}}{n^0_b}\right) - 1$ ensures that $n_{ub}$ and $n_b$ relax toward their respective steady-state values. These switching probabilities depend on the instantaneous population of nucleosomes in each state and collectively ensure the maintenance of a steady state [7, 8].

Equations S8 and S9 apply when $n_{ub} \geq n_b$. When $n_b > n_{ub}$, Eq. S8 is modified by replacing the denominator constant 1 with $\eta$, where $\eta = 2\left(\frac{n^0_b}{n^0_{ub}}\right) - 1$, and in Eq. S9, $\eta$ is replaced by 1, and the asymmetry parameter becomes $A_0 = n^0_b - n^0_{ub}$ [7, 8].

Simulations are performed at average remodeler-bound fractions $\langle n_b/n_N\rangle = 0.1$ and $0.2$, with a total nucleosome count of $n_N = 1296$. The parameter $\mu = 1$ was chosen to ensure small fluctuations between the remodeler-bound and remodeler-unbound nucleosome populations.

The number $n_{\rm sw}$ controls the frequency of nucleosome state-switching attempts and therefore modulates the average residence time of a nucleosome in each state before switching. We define the switching time as

$$t_{\rm a} = \left(\frac{n_b}{n_{\rm sw}}\right)\tau,$$

which sets the characteristic time scale over which $n_b$ nucleosomes are considered for state switching. We perform simulations with $n_{\rm sw} = 0, 0.05n_b, 0.1n_b, 0.2n_b, 0.5n_b, n_b$, corresponding to switching times $t_{\rm a} = \infty$, $20\tau$, $10\tau$, $5\tau$, $2\tau$, $\tau$, respectively. The case $n_{\rm sw} = 0$ corresponds to the passive limit, with $t_{\rm a} \to \infty$.

Unless otherwise specified, simulations of the single-bead nucleosome model contain $N_{fiber} = 108$ chromatin fibers.

### B. Active Chromatin with Hydrodynamic Interactions

To examine how forces generated by chromatin remodelers influence condensate dynamics via hydrodynamic coupling, we perform simulations with explicit solvent using Dissipative Particle Dynamics (DPD) [9, 10]. We consider two scenarios: (a) remodeler binding modifies only the effective nucleosome–nucleosome interactions (as described in Section I A); and (b) in addition to altering these interactions, remodelers generate ATP-driven mechanical activity, which, to leading order, is represented as a force dipole (see Fig. 1(d)).

In the DPD model, the solvent and chromatin are represented as coarse-grained beads of diameter $\sigma$. There are four types of beads representing: (i) solvent, (ii) linker DNA, (iii) nucleosomes, and (iv) remodeler-bound nucleosomes. The non-bonded interactions between all beads are soft repulsive. The detailed forms of these interactions are given below.

The equations of motion governing the system dynamics are defined as follows. The position $\vec{r}_i$ of bead $i$ with mass $m_i$ evolves according to Newton's equation of motion,

$$m_i \frac{d\vec{v}_i}{dt} = \sum_{j \neq i} \left( \vec{F}_{ij}^C + \vec{F}_{ij}^D + \vec{F}_{ij}^R \right) + \vec{F}_i^A, \qquad \text{(S10)}$$

where $\vec{v}_i$ is the velocity of bead $i$. The terms $\vec{F}_{ij}^C$, $\vec{F}_{ij}^D$, and $\vec{F}_{ij}^R$ represent the conservative, dissipative, and random DPD forces acting between beads $i$ and $j$, while $\vec{F}_i^A$ denotes the active force associated with remodeler activity. The explicit forms of these forces are defined below.

In this coarse-grained model, length, energy, and time are measured in units of the bead diameter $\sigma$, $k_{\mathrm{B}}T$, and the characteristic timescale $\tau = \sigma\sqrt{m/k_{\mathrm{B}}T}$, respectively. For mapping to physical units, $\sigma$ corresponds to 12.5 bp, and $\tau \approx 4 \times 10^{-10}$ s.

The conservative force acting on bead $i$ is given by

$$\vec{F}_i^C = \sum_{j \neq i} \vec{F}_{ij}^{\mathrm{nb}} - \sum_{j = i \pm 1} \nabla_i U_{ij}^{\mathrm{b}} - \nabla_i U_i^{\mathrm{bend}}, \qquad \text{(S11)}$$

where the bending potential $U_i^{\mathrm{bend}}$ acts on the bead triplet $(i-1, i, i+1)$ along the chain.

The non-bonded interaction is given by the standard DPD soft repulsion,

$$\vec{F}_{ij}^{\mathrm{nb}} = a_{ij} \left( 1 - \frac{r_{ij}}{r_c} \right) \hat{r}_{ij}, \qquad r_{ij} < r_c, \qquad \text{(S12)}$$

and zero otherwise. Here the cutoff distance is $r_c = 1$. The repulsion parameter $a_{ij}$ controls the effective interaction between bead types. For nucleosome–nucleosome pairs, with no remodeler bound to it, $a_{ij} = 14$, which promotes condensate formation, whereas for all other non-bonded bead pairs $a_{ij} = 50$.

Bonded interactions ($U_{ij}^{\mathrm{b}}$) are modeled using harmonic springs [Eq. S3], with parameters $k_b = 200$ and $r_0 = 0.87$.

Fiber stiffness is modeled using a bending potential acting on triplets of beads (Eq. S7), with $\kappa = 22$ for linker-linker-linker triplets and preferred angle $\theta_0 = 180°$. For all other triplets we set $\kappa = 0$. These parameters reproduce the effective persistence length of DNA in our coarse-grained model [5, 6].

The dissipative and random forces implement a pairwise momentum-conserving thermostat and are given by

$$\vec{F}_{ij}^D = -\gamma_{ij} w^2(r_{ij}) (\hat{r}_{ij} \cdot \vec{v}_{ij}) \hat{r}_{ij}, \qquad \text{(S13)}$$

$$\vec{F}_{ij}^R = \sigma_{ij} w(r_{ij}) \xi_{ij} \hat{r}_{ij}, \qquad \text{(S14)}$$

where $\vec{v}_{ij} = \vec{v}_i - \vec{v}_j$ is the relative velocity and $\xi_{ij}$ is a Gaussian random variable with zero mean and unit variance. The weight function is defined as

$$w(r_{ij}) = 1 - \frac{r_{ij}}{r_c}, \qquad r_{ij} < r_c,$$

and $w(r_{ij}) = 0$ otherwise. Because both $\vec{F}_{ij}^D$ and $\vec{F}_{ij}^R$ are pairwise antisymmetric and depend only on relative coordinates, momentum is conserved locally, enabling the emergence of hydrodynamic interactions [10]. To satisfy the fluctuation–dissipation theorem and maintain the system at temperature $T$, the noise amplitude satisfies $\sigma_{ij}^2 = 2\gamma_{ij} k_{\mathrm{B}} T$. Unless otherwise specified, we set $\gamma_{ij} = 9$. Together with the chosen conservative interaction parameters $a_{ij}$, this yields a Schmidt number Sc $\approx 2.58$, ensuring that momentum diffusion dominates over mass diffusion and that the solvent exhibits liquid-like hydrodynamic behavior (see Sec. II 1 in SI) [11, 12].

The active force $\vec{F}_i^A$ represents a force dipole aligned with the bond connecting the remodeler-bound nucleosome bead and its bonded linker bead. The dipole is implemented through two spatially separated force distributions defined within spherical regions of radius $R_a = 1.3$ centered on the nucleosome bead $i$ and its bonded linker bead $j$ ($j = i \pm 1$), as illustrated in the inset of Fig. 1d. Beads within these two regions experience equal and opposite forces along the unit bond vector $\hat{r}_{ij} = (\vec{r}_j - \vec{r}_i)/|\vec{r}_j - \vec{r}_i|$. The force is distributed among the beads in these volumes such that the total force within the nucleosome-centered sphere equals $-F_0 \hat{r}_{ij}$, while that within the linker-centered sphere equals $+F_0\hat{r}_{ij}$, ensuring overall force balance. The active force acting on a bead $k$ within the nucleosome-centered sphere ($r_{ik} < R_a$) is therefore given by

$$\vec{F}_k^A = -F_0 \frac{\mathcal{W}(r_{ik})}{\sum_{k' \in \mathcal{S}_i} \mathcal{W}(r_{ik'})} \hat{r}_{ij}, \tag{S15}$$

while for beads within the linker-centered sphere ($r_{jk} < R_a$),

$$\vec{F}_k^A = +F_0 \frac{\mathcal{W}(r_{jk})}{\sum_{k' \in \mathcal{S}_j} \mathcal{W}(r_{jk'})} \hat{r}_{ij}. \tag{S16}$$

$\mathcal{S}_i = \{k \notin \{i-1, i, i+1\} \mid r_{ik} < R_a\}$ denotes the set of beads within the spherical region of radius $R_a$ centered at bead $i$, excluding bead $i$ and its bonded neighbors ($i \pm 1$). The weight function is defined as

$$\mathcal{W}(r_{ik}) = \left(1 - \frac{r_{ik}}{R_a}\right)^2, \qquad r_{ik} < R_a,$$

and zero otherwise. The normalization ensures that the total magnitude of the force distributed within each sphere equals $F_0$. By construction, $\sum_k \vec{F}_k^A = 0$, ensuring overall force balance. With the sign convention used in Eqs. (S15) and (S16), $F_0 > 0$ corresponds to an extensile dipole, in which forces are directed outward along the nucleosome–linker bond. A contractile dipole is obtained by reversing the force directions, equivalently by taking $F_0 < 0$ (see inset, Fig. 1d).

We simulate $1.92 \times 10^5$ beads in total, corresponding to a number density of 3 [10]. The system contains $N_{fiber} = 20$ chromatin fibers, each comprising 150 beads. Nucleosomes are placed at uniformly spaced intervals of four linker beads along each fiber, yielding $n_N = 600$ nucleosomes. The remaining beads serve as explicit solvent.

Periodic boundary conditions are applied in all three spatial directions to mimic an unbounded fluid and to allow momentum to propagate across the simulation box. Simulations are performed at a fixed average remodeler-bound fraction $\langle n_b/n_N \rangle = 0.1$. We use force dipole magnitudes $F_0 = 0$, 30, 40, corresponding to extensile activity ($F_0 > 0$). Contractile dipoles are obtained by taking $F_0 < 0$. These magnitudes are comparable to experimentally reported forces for the RSC remodeler [13].

### II. SIMULATION DETAILS AND ANALYSIS

All simulations were performed using the LAMMPS molecular dynamics package [14, 15] (version 29 Aug 2024), with GPU acceleration and custom fixes. Dissipative particle dynamics simulations employed the Shardlow integration scheme [16–18].

#### 1. *Schmidt number*

The Schmidt number is defined as the ratio of the kinematic viscosity ($\nu$) to the diffusion coefficient ($D$),

$$\mathrm{Sc} = \frac{\nu}{D}. \tag{S17}$$

Since $\nu = \eta/\rho$, where $\eta$ is the shear viscosity and $\rho$ is the number density, this can be written as

$$\mathrm{Sc} = \frac{\eta}{\rho D}. \tag{S18}$$

To estimate Sc, we simulate a system consisting only of solvent beads at a number density $\rho = 3$ in a cubic box of size $L = 30$, with conservative interaction parameter $a_{ij} = 50$.

The shear viscosity $\eta$ is calculated using the Green–Kubo relation [12],

$$\eta = \frac{V}{k_\mathrm{B}T} \int_0^\infty \langle P_{\alpha\beta}(0)\, P_{\alpha\beta}(t) \rangle \, dt, \tag{S19}$$

where $V$ is the system volume and $P_{\alpha\beta}$ ($\alpha\beta = xy, xz, yz$) are the off-diagonal components of the pressure tensor. The time integral is evaluated numerically using the trapezoidal rule,

$$\eta(t_n) = \frac{V}{k_\mathrm{B}T} \sum_{i=1}^{n} \frac{C(t_i) + C(t_{i-1})}{2} (t_i - t_{i-1}), \tag{S20}$$

where $C(t) = \langle P_{\alpha\beta}(0) P_{\alpha\beta}(t) \rangle$ is the stress autocorrelation function. We obtain $\eta \approx 1.32$

The diffusion coefficient $D$ of beads is obtained from the long-time diffusive regime of the mean-squared displacement, $\langle \Delta r^2(t) \rangle = 6Dt$, yielding $D = 0.1703$.

Using these values, we get $\mathrm{Sc} \approx 2.58$.

#### 2. *Estimation of Inter-nucleosome Interaction Strength Using FRAP Timescales*

The relevant inter-nucleosome interaction strength is identified by comparing the characteristic bead displacement in simulations over the experimental FRAP recovery time with the condensate length scale probed experimentally. The bead MSD is fitted to $\langle \Delta r^2(t) \rangle = 6Dt^\alpha$, where $D$ is the generalized diffusion coefficient and $\alpha$ is the dynamic exponent. The fitted diffusion coefficient, $D_{\rm sim}$ $(\sigma^2/\tau^\alpha)$, is converted to physical units, $D_{\rm phys}$ $(\mu{\rm m}^2/{\rm s}^\alpha)$. Using the experimental FRAP recovery time, $t \approx 5$ min, the characteristic bead displacement in the simulation is estimated as $\Delta r = \sqrt{6 D_{\rm phys} t^\alpha}$.

**TABLE S1:** Generalized diffusion coefficient, dynamic exponent, and estimated bead displacement for different inter-nucleosome interaction strengths $\epsilon_{\rm NN}$.

| $\boldsymbol{\epsilon}_{\rm NN}(k_{\rm B}T)$ | $D_{\rm sim}$ $(\sigma^2/\tau^\alpha)$ | $\boldsymbol{\alpha}$ | $D_{\rm phys}$ $(\mu{\rm m}^2/{\rm s}^\alpha)$ | $\Delta r$ ($\mu$m) |
|---|---|---|---|---|
| 3 | 0.0562 | 0.54 | 0.134 | 4.27 |
| 4 | 0.0019 | 0.76 | 0.516 | 15.37 |
| 5 | 0.0004 | 0.85 | 0.948 | 27.47 |
| 6 | 0.0006 | 0.80 | 0.389 | 14.91 |

#### 3. *Mixing parameter*

In experiments, fluidity is typically assessed using FRAP, where a portion of a fluorescent condensate is bleached and the fluorescence recovery is monitored over time [1, 2]. Following this idea, we tag beads in one half of the condensate. At each time $t$, we compute the pairwise distance $d(t)$ between all beads, and the distance $d'(t)$ between tagged and untagged beads. These values are binned. The mixing function $M(t)$ is then defined as

$$M(t) = \sum_i \frac{(d'_i - d_i)^2}{d_i}, \tag{S21}$$

where the index $i$ is summed over all bins. We normalize $M$ to lie between 0 and 1, and define the mixing parameter as $m(t) = 1 - M(t)$, where $m(t) = 0$ corresponds to complete separation between tagged and untagged beads, and $m(t) = 1$ corresponds to their complete intermixing [19].

#### 4. *Pair-correlation function*

The pair-correlation function is defined as [12]

$$g_N(r) = \frac{1}{4\pi r^2 \rho n_N} \Big\langle \sum_{i=1}^{n_N} \sum_{j\neq i}^{n_N} \delta(r - r_{N,ij}) \Big\rangle, \tag{S22}$$

where $\rho$ is the average nucleosome density in the condensate.

#### 5. *Radius of gyration*

The radius of gyration is defined as

$$R_g = \sqrt{\frac{1}{n} \Big\langle \sum_{i=1}^{n} |\vec{r}_i - \vec{R}_{cm}|^2 \Big\rangle}. \tag{S23}$$

#### 6. *Radial bead density*

Radial bead density is calculated by binning bead positions into concentric spherical shells of thickness $\Delta r$ centered on the condensate center of mass. For fiber type $\nu$, the radial bead density is given by

$$\rho_\nu(r) = \frac{N_\nu(r)}{V(r)}, \tag{S24}$$

where $N_\nu(r)$ is the number of beads belonging to fiber type $\nu$ in the shell at radius $r$, and $V(r) = 4\pi \left[(r+\Delta r)^3 - r^3\right]/3$ is the corresponding shell volume.

#### 7. *Linking number:*

The Gaussian linking number $Lk_G$ for two curves, $C_1$ and $C_2$, is defined as

$$Lk_G = \frac{1}{4\pi} \int_{C_1} \int_{C_2} (d\vec{r}_1 \times d\vec{r}_2) \cdot \frac{\vec{r}_2 - \vec{r}_1}{|\vec{r}_2 - \vec{r}_1|^3}, \tag{S25}$$

where $\vec{r}_1$ and $\vec{r}_2$ are points on curves $C_1$ and $C_2$, respectively [20]. For fibers discretized into $N_1$ and $N_2$ beads with positions $\vec{r}_1^{\,i}$ and $\vec{r}_2^{\,j}$, the Gaussian linking number can be approximated as

$$Lk_G = \frac{1}{4\pi} \sum_{i=1}^{N_1-1} \sum_{j=1}^{N_2-1} \left(\Delta\vec{r}_1^{\,i} \times \Delta\vec{r}_2^{\,j}\right) \cdot \frac{\vec{r}_2^{\,j} - \vec{r}_1^{\,i}}{|\vec{r}_2^{\,j} - \vec{r}_1^{\,i}|^3}, \tag{S26}$$

where $\Delta\vec{r}_1^{\,i} = \vec{r}_1^{\,i+1} - \vec{r}_1^{\,i}$ and $\Delta\vec{r}_2^{\,j} = \vec{r}_2^{\,j+1} - \vec{r}_2^{\,j}$ represent tangent vectors at segments along the two fiber chains. Since the fibers considered here are open linear chains, $Lk_G$ is generally non-integer and is used only as a geometric measure of inter-fiber linking/entanglement. Pairs with $|Lk_G| < 0.5$ are classified as weakly linked and are not counted as entangled.

### III. FIGURES

**(a)** **(b)**

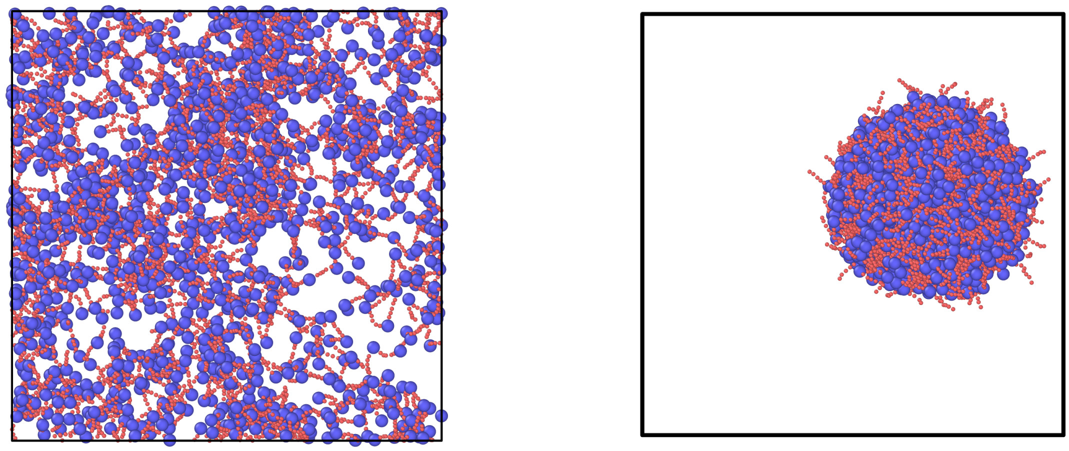

**FIG. S1:** Snapshots of chromatin fibers at different nucleosome–nucleosome interaction strengths ($\epsilon_{\rm NN}$), showing condensate formation for $\epsilon_{\rm NN} \gtrapprox 3k_{\rm B}T$: **(a)** $\epsilon_{\rm NN} = 1k_{\rm B}T$ and **(b)** $\epsilon_{\rm NN} = 3k_{\rm B}T$. Blue beads represent nucleosomes, and red beads represent linkers.

**(a)** **(b)**

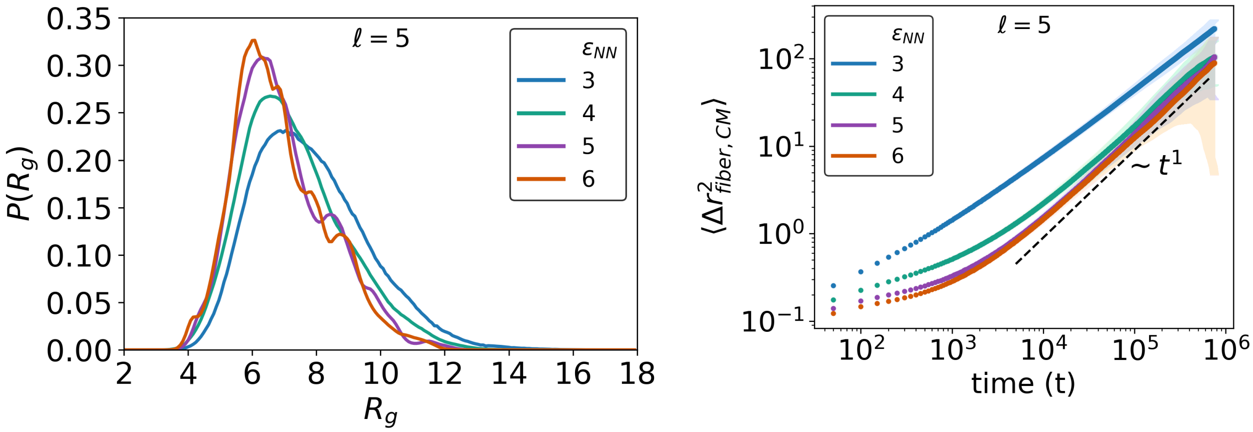


**FIG. S2:** **(a)** Probability distribution of the fiber radius of gyration, $P(R_g)$, for different inter-nucleosome interaction strengths $\epsilon_{\rm NN}$ at fixed linker length $\ell = 5$. **(b)** MSD of the fiber center of mass (CM) relative to the condensate CM for $\ell = 5$ and different inter-nucleosome interaction strengths $\epsilon_{\rm NN}$.

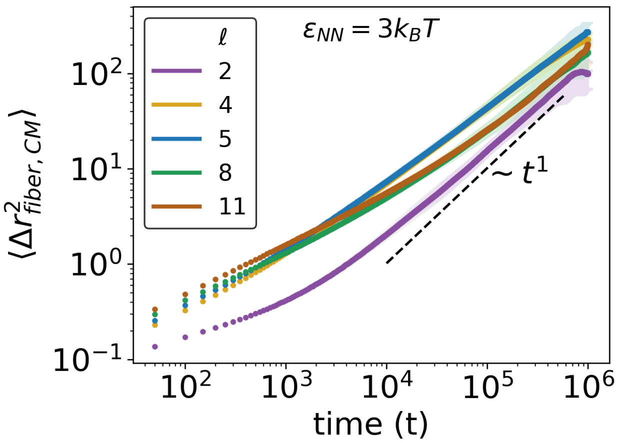


**FIG. S3:** MSD of the fiber CM with respect to the condensate CM shown for $\epsilon_{\rm NN} = 3\,k_{\rm B}T$ and various linker lengths $\ell$.

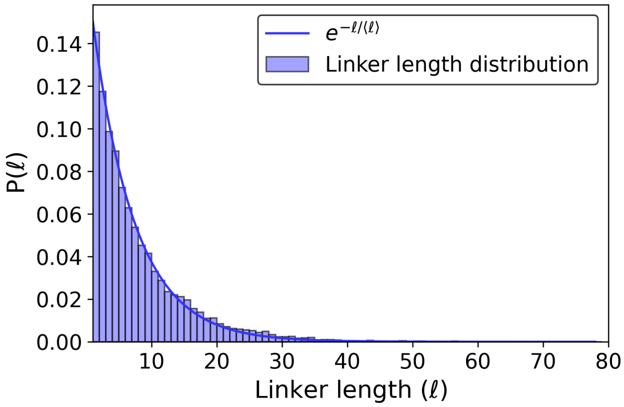


**FIG. S4:** Distribution of linker lengths for irregular fibers, where the number of nucleosomes per fiber is sampled from a Gaussian distribution; see Results section II 3. The average linker length is $\langle \ell \rangle = 6.4$.

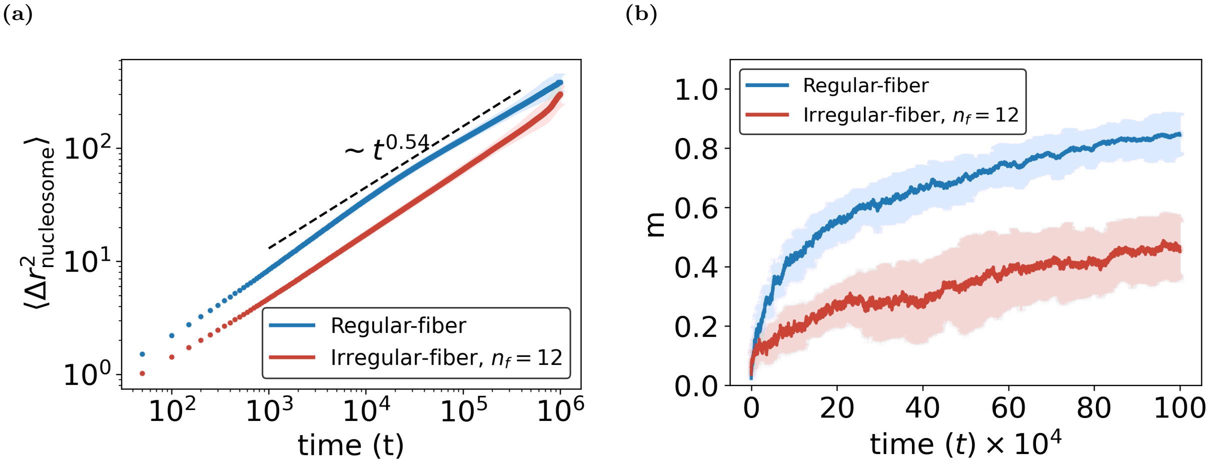


**FIG. S5: (a)**Nucleosome MSD in condensates formed from regular and irregular fibers with twelve nucleosomes per fiber. **(b)** Mixing parameter over time for the same systems as in (a).

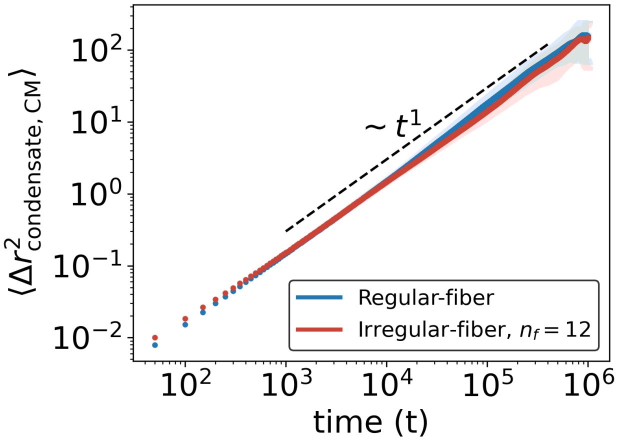


**FIG. S6:** Condensate CM MSD for condensates formed from regular and irregular fibers with twelve nucleosomes per fiber.

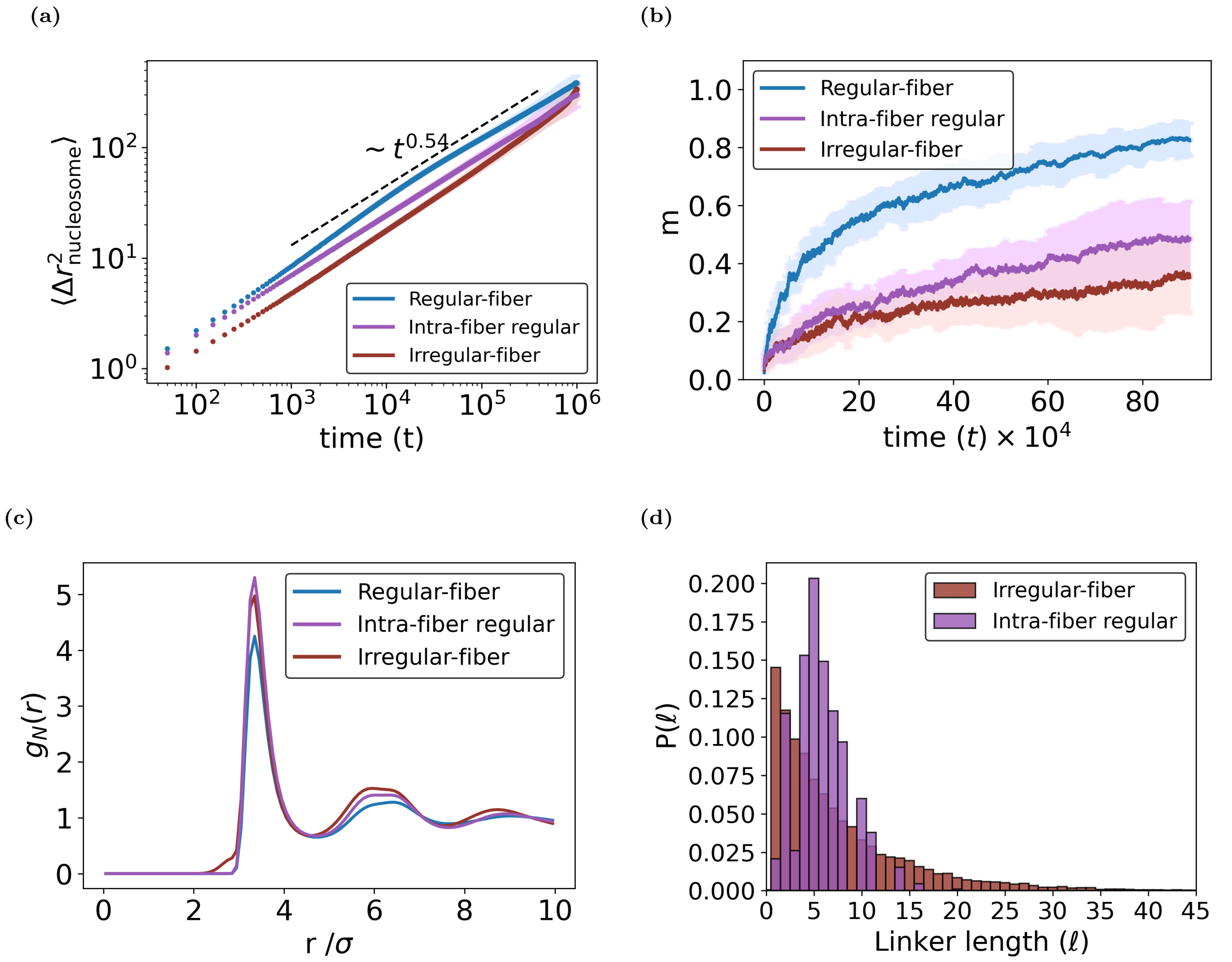


**FIG. S7:** Comparison of condensate properties for the intra-fiber regular system. **(a)** MSD of nucleosomes in the condensate center-of-mass frame. **(b)** Mixing parameter as a function of time. **(c)** Pair correlation function $g_N(r)$ of nucleosomes. **(d)** Linker length distributions for the intra-fiber regular and irregular-fibers.

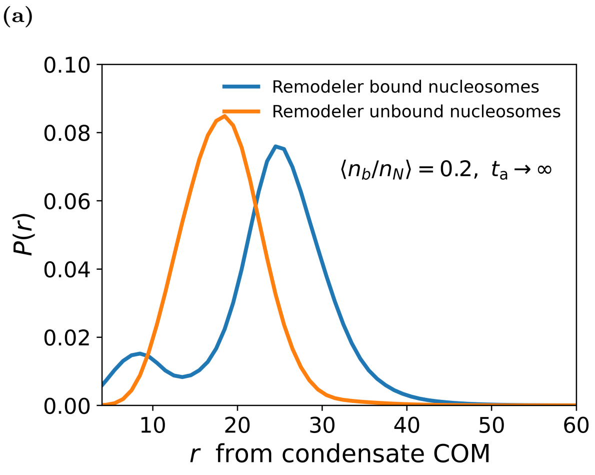


**FIG. S8:** Normalized radial probability distributions of remodeler-bound and remodeler-unbound nucleosomes measured from the condensate CM. **(a)** Passive case with $\langle n_b/n_N \rangle = 0.2$ and $t_\mathrm{a} \to \infty$. **(b)** Nucleosome state switching with $\langle n_b/n_N \rangle = 0.2$ and $t_\mathrm{a} = \tau$. The distributions are obtained by binning nucleosomes according to their radial distance from the condensate CM and normalizing each profile by the number of nucleosomes of the corresponding type.

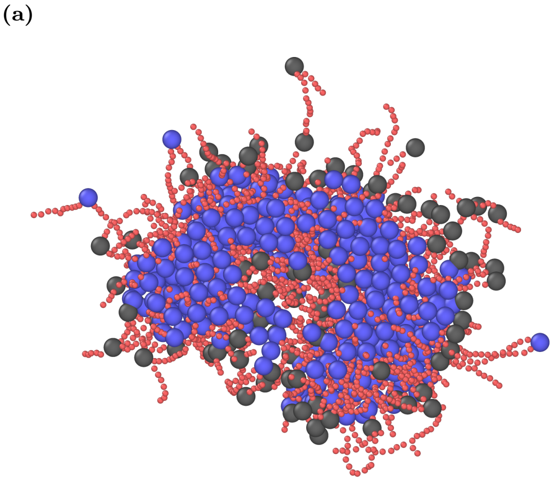


**FIG. S9:** Slab-cut snapshots of condensates showing the spatial distribution of remodeler-bound nucleosomes (grey) and remodeler-unbound nucleosomes (blue). **(a)** Passive case with $\langle n_b/n_N \rangle = 0.2$. **(b)** Nucleosome state switching with $\langle n_b/n_N \rangle = 0.2$ and $t_\mathrm{a} = \tau$. The slab is taken along the $z$ direction with a width of 15.

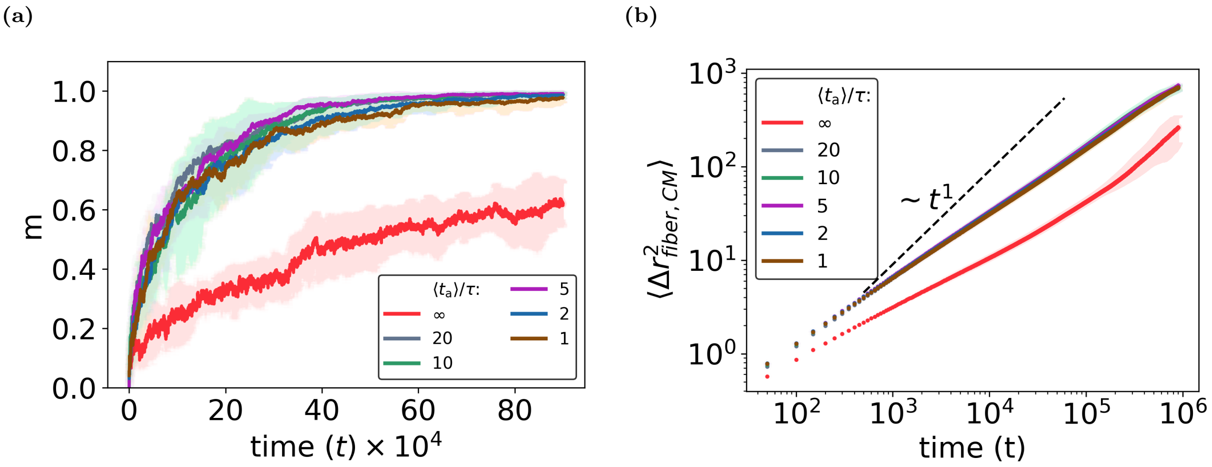


**FIG. S10:** **(a)** Mixing of beads as a function of time for $\langle n_b/n_N \rangle = 0.2$ at different $t_a$. **(b)** MSD of the fiber center of mass (CM) in the condensate CM frame for $\langle n_b/n_N \rangle = 0.2$.

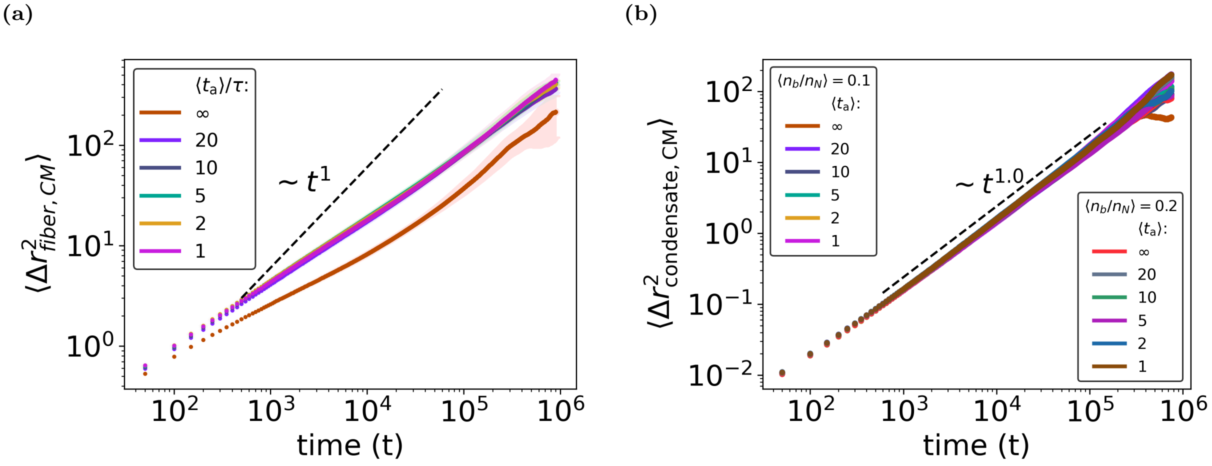


**FIG. S11:** **(a)** MSD of the fiber CM in the condensate CM frame for $\langle n_b/n_N \rangle = 0.1$. **(b)** MSD of the condensate CM for different $\langle n_b/n_N \rangle$ and switching times $t_a$.

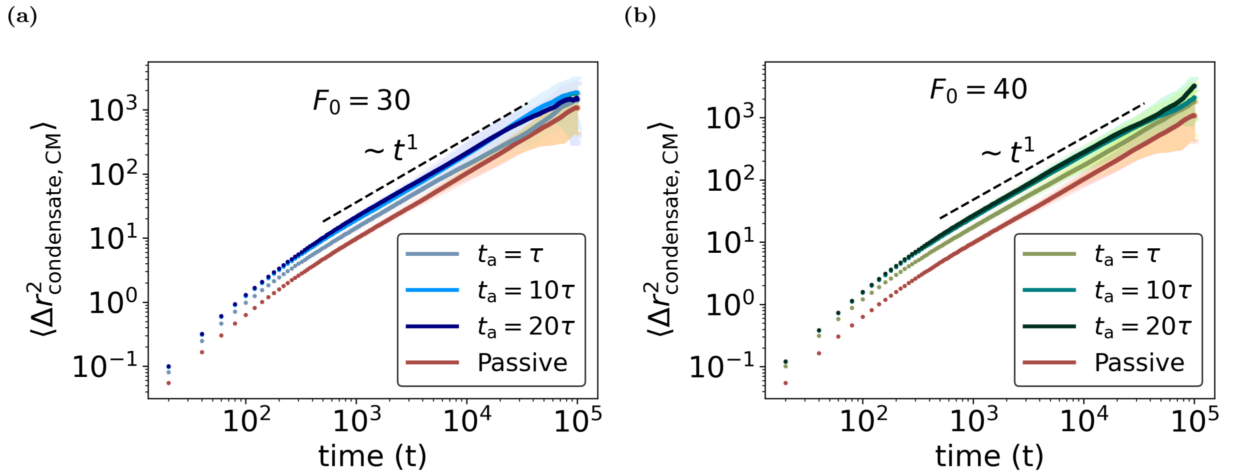


**FIG. S12:** MSD of the condensate center of mass for extensile dipolar forces with **(a)** $F_0 = 30$ and **(b)** $F_0 = 40$, for varying $t_{\mathrm{a}}$. The passive case corresponds to $F_0 = 0$ and $t_{\mathrm{a}} = \infty$. Shaded regions represent the standard deviation.

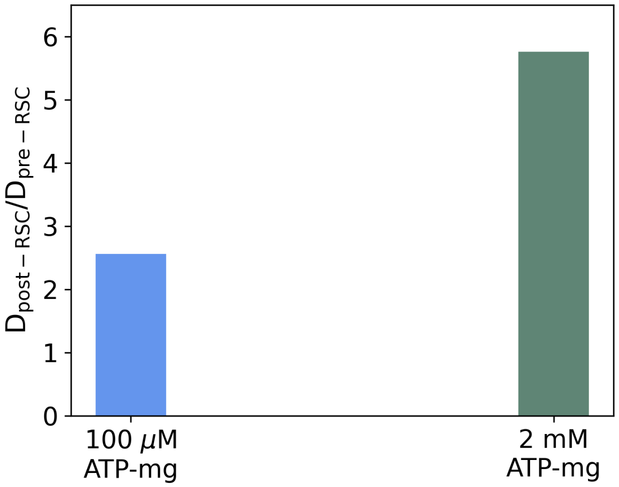


**FIG. S13:** Ratio of condensate diffusion coefficient after and before RSC remodeler activity, $D_{\text{post-RSC}}/D_{\text{pre-RSC}}$, extracted from experimental data reported by Moore *et al.* [2]. The effective diffusion coefficient is estimated from the reported condensate displacement over a time of 20 s, assuming diffusive motion, using $D = \langle \Delta r^2 \rangle / (4t)$.

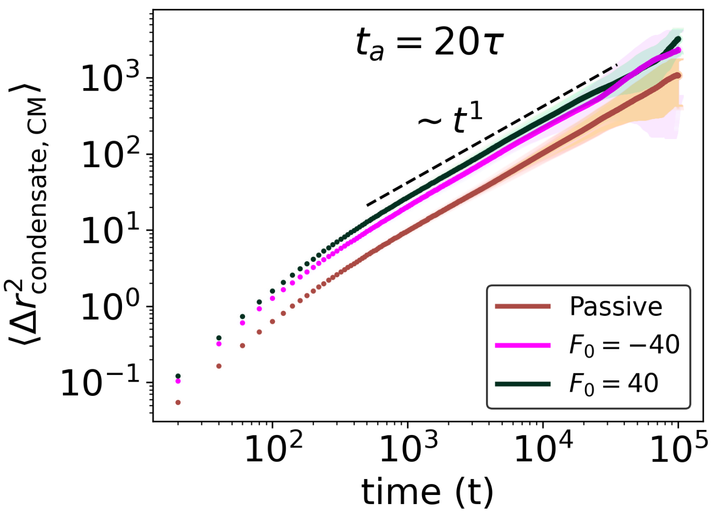


**FIG. S14:** MSD of the condensate CM for different dipolar activities at $t_a = 20\tau$. Positive $F_0$ denotes extensile dipolar forces, while negative $F_0$ ($F_0 = -40$) corresponds to contractile dipolar forces.

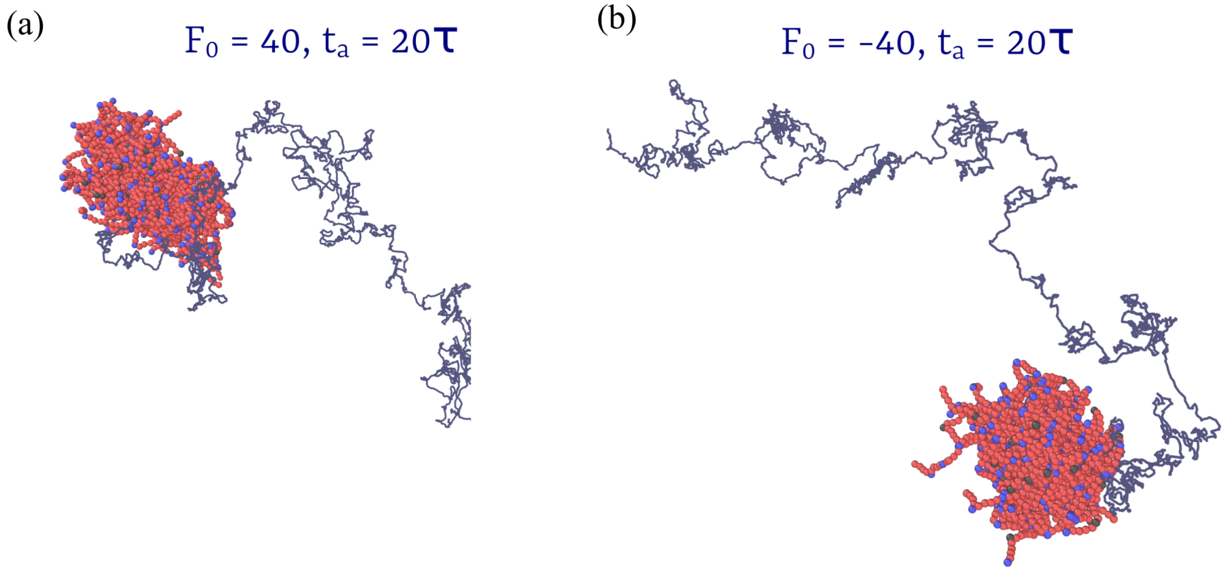


**FIG. S15:** Snapshots of center-of-mass trajectories of condensates. **(a)** Extensile force dipole, $F_0 = 40$, $t_a = 20\tau$, and **(b)** contractile force dipole, $F_0 = -40$, $t_a = 20\tau$. The grey lines show the paths traced by the condensate center of mass over a fixed duration of $10^5\tau$.

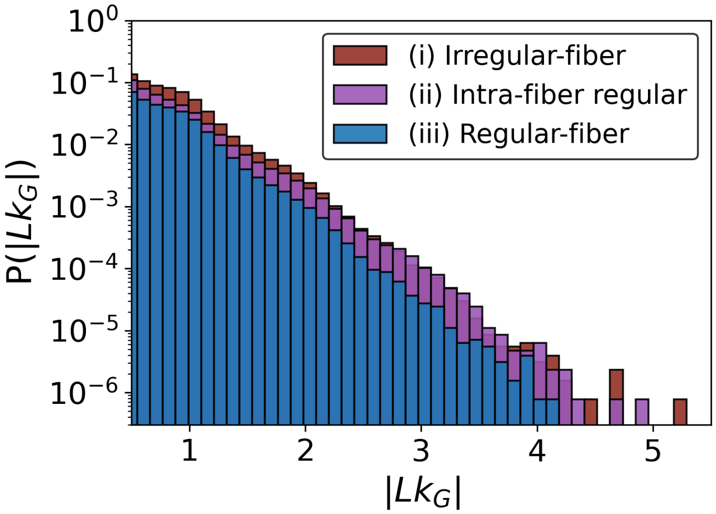


**FIG. S16:** Probability distribution of the Gaussian linking number between fiber pairs in the condensate composed of (i) Irregular-fiber (ii) Intra-fiber regular, and (iii) Regular-fiber.

### IV. MOVIES

**Movie S1. Active dipolar forces resulting from the action of nucleosome remodelers enhance condensate center-of-mass motion via hydrodynamic interactions.** Representative trajectories of chromatin condensates are shown for different active dipolar force strengths, $F_0$, and nucleosome state-switching times, $t_{\rm a}$. The movie compares the passive case, $F_0 = 0$ and $t_{\rm a} \to \infty$, with nucleosome state switching only, $F_0 = 0$ and $t_{\rm a} = \tau$, and extensile dipolar activity, $F_0 = 30$ and 40 with $t_{\rm a} = 10\tau$. The condensate CM MSD for these cases is shown in Fig. 6b.

**Movie S2. Condensate center-of-mass motion is independent of the sign of the force dipole.** Representative trajectories compare extensile ($F_0 = 40$) and contractile ($F_0 = -40$) dipolar activity at fixed nucleosome state-switching time, $t_{\rm a} = 20\tau$. The corresponding condensate CM MSD is shown in Fig. S14.

[1] B. A. Gibson, L. K. Doolittle, M. W. G. Schneider, L. E. Jensen, N. I. Gamarra, L. Henry, D. W. Gerlich, S. Redding, and M. K. Rosen, Cell **179**, 470 (2019).
[2] C. Moore, E. Wong, U. Kaur, U. S. Chio, Z. Zhou, M. Ostrowski, K. Wu, I. Irkliyenko, S. Wang, V. Ramani, and G. J. Narlikar, Science **390**, eadr0018 (2025).
[3] S. Kadam, K. Kumari, V. Manivannan, S. Dutta, M. K. Mitra, and R. Padinhateeri, Nature Communications **14**, 10.1038/s41467-023-39832-3 (2023).
[4] A. J. Beel, M. Azubel, P.-J. Matteï, and R. D. Kornberg, Molecular Cell **81**, 4369 (2021).
[5] P. J. Hagerman, Annual Review of Biophysics and Biophysical Chemistry **17**, 265 (1988).
[6] C. G. Baumann, S. B. Smith, V. A. Bloomfield, and C. Bustamante, Proceedings of the National Academy of Sciences of the United States of America **94**, 6185 (1997).
[7] S. Sahoo, S. Kadam, R. Padinhateeri, and P. B. Sunil Kumar, Soft Matter **20**, 4621 (2024).
[8] W. Im, S. Seefeld, and B. Roux, Biophysical Journal **79**, 788 (2000).
[9] P. Español and P. Warren, Europhysics Letters **30**, 191 (1995).
[10] R. D. Groot and P. B. Warren, The Journal of Chemical Physics **107**, 4423 (1997).
[11] R. B. Bird, W. E. Stewart, and E. N. Lightfoot, *Transport Phenomena*, 2nd ed. (Wiley, 2002).
[12] J.-P. Hansen and I. R. McDonald, *Theory of Simple Liquids*, 3rd ed. (Academic Press, London, 2006).
[13] G. Sirinakis, C. R. Clapier, Y. Gao, R. Viswanathan, B. R. Cairns, and Y. Zhang, The EMBO Journal **30**, 2364 (2011).
[14] A. P. Thompson, H. M. Aktulga, R. Berger, D. S. Bolintineanu, W. M. Brown, P. S. Crozier, P. J. in 't Veld, A. Kohlmeyer, S. G. Moore, T. D. Nguyen, R. Shan, M. J. Stevens, J. Tranchida, C. Trott, and S. J. Plimpton, Computer Physics Communications **271**, 108171 (2022).
[15] W. M. Brown, P. Wang, S. J. Plimpton, and A. N. Tharrington, Comput. Phys. Commun. **182**, 898 (2011).
[16] J. R. Gissinger, I. Nikiforov, Y. Afshar, B. Waters, M.-k. Choi, D. S. Karls, A. Stukowski, W. Im, H. Heinz, A. Kohlmeyer, and E. B. Tadmor, Journal of Physical Chemistry B **128**, 3282 (2024).
[17] J. P. Larentzos, J. K. Brennan, J. D. Moore, M. Lísal, and W. D. Mattson, Computer Physics Communications **185**, 1987 (2014).
[18] M. Lísal, J. K. Brennan, and J. Bonet Avalos, Journal of Chemical Physics **135**, 204105 (2011).
[19] P. Vizjak, D. Kamp, N. Hepp, A. Scacchetti, M. G. Pisfil, J. Bartho, M. Halic, P. B. Becker, M. Smolle, J. Stigler, and F. Mueller-Planitz, Nature Structural & Molecular Biology **31**, 1331 (2024).
[20] R. Ahmad, S. Paul, and S. Basu, Physical Review E **101**, 022503 (2020).
